# Expression of four mitochondrial tRNAs from only two loci

**DOI:** 10.1101/2025.11.12.688142

**Authors:** Jessica M. Warren, Kasavajhala V.S.K. Prasad, Anistynn M. Mendez, Stephanie Temnyk, John P. McCutcheon

## Abstract

Transfer RNAs (tRNAs) are among the few genes retained in animal mitochondrial genomes after more than a billion years of gene loss. These ancient bacterial vestiges are often structurally aberrant and less stable than their bacterial or cytosolic tRNA counterparts. In some lineages, mitochondrial tRNAs (mt-tRNAs) have become so truncated that the loss of one or both arms has expanded our understanding of what constitutes a functional tRNA. Here, we report another radical departure from canonical tRNA gene architecture: two overlapping tRNAs produced from opposite strands of the same locus. These ‘mirror’ tRNA pairs eliminate the need to retain separate loci for all tRNA genes, as a single locus can produce tRNAs to decode two different amino acids. We show that these mirror tRNAs are aminoacylated and demonstrate their presence in mitoribosomes. Furthermore, mirror tRNAs display strand-specific patterns of nucleotide modification and RNA editing, reflecting specific post-transcriptional maturation that depends on transcriptional orientation. To our knowledge, this demonstration of functional, bidirectional tRNA expression is a first for any genome or organism and reveals an unexpected strategy by which mitochondrial genomes maintain a complete set of tRNAs in the face of unrelenting gene loss. The discovery of mirror tRNAs has broad implications for the evolution of tRNA-interacting enzymes, mitochondrial biology, and even the origins of the protein synthesis machinery itself.

## Introduction

The mitochondrial genomes of most bilaterian animals have been reduced to a remarkably conserved set of the same 37 genes, 22 of which encode transfer RNAs (1). These 22 mitochondrial tRNAs (mt-tRNAs) are considered a minimally sufficient set of tRNAs to decode all 60 codons for the 20 amino acids in the mitochondrial genetic code (2). This almost one-to-one relationship between amino acids and tRNAs is in striking contrast to the more than 50 species of tRNAs used for decoding all codons in prokaryotic or eukaryotic cytosolic translation and represents a fascinating example of a near-minimal protein synthesis system (3). However, this stubborn retention of mt-tRNAs in animals is not universal in mitochondrial evolution. The mitochondrial genomes (mitogenomes) of most non-animal eukaryotes have lost tRNAs beyond this minimal set and rely on the import of nuclear-encoded tRNAs to maintain mitochondrial translation (4, 5). Therefore, encoding a full set of native mt-tRNAs, once thought to be the general rule, actually represents an exceptional case of genetic entrenchment.

This conservation of tRNAs on the animal mitogenome has had profound consequences for the structural evolution of mt-tRNAs. Bilaterian mitogenomes experience a much higher rate of sequence evolution than the nuclear genome due to their high mutation rates, asexual reproduction, limited recombination, and frequent population bottlenecks (6). Over 500 million years of animal evolution, the accumulation of slightly deleterious mutations has resulted in animal mt-tRNAs being shorter and having overall less thermal stability than their nuclear counterparts(7–9). tRNA truncation has been so extensive in some lineages that numerous mt-tRNAs have lost the entire D- or T-arm, and in the most extreme cases, both arms have been lost and the mt-tRNA is now just a simple stem-loop structure (10–12).

Here, we report a new way in which mt-tRNA evolution has pushed the bounds of genetic compaction. We show that in the mitochondrion of the citrus mealybug (*Planococcus citri*), four functional tRNAs are produced from two tRNA loci, and that the expression, modification, and aminoacylation of these four tRNAs provides *P. citri* mitochondria with its complete set of 22 mt-tRNAs from only 20 mt-tRNA loci.

## Results

### The *P. citri* mitogenome is the expected size for an animal but is extremely A/T biased

A previous *P. citri* mitogenome has been published (GenBank: OX465514) as part of the Wellcome Sanger Tree of Life Programme. However, we noticed that this genome was almost twice the size (at 30,900 bp) of other assembled Coccoidea (scale insects) mitogenomes, including the closely related *Phenacoccus manihoti* (14,965 bp). The published *P. citri* mitogenome assembly also appears to contain a large, inverted duplication. We suspected that these features were due to misassembly errors. We therefore re-sequenced and re-assembled the *P. citri* mitogenome using Oxford Nanopore on isolated mitochondrial DNA. This new assembly resulted in a 14,910 bp circular molecule that shares 98.8% sequence identity with the previously published mitogenome. The overall base pair composition of the mitogenome is extremely A/T biased with 90.8% A/T (9.2% G/C) and contains a control region with a large “ATAT” tandem duplication. This new *P. citri* mitogenome assembly was deposited on GenBank: PZ166947.

### Multiple *P. citri* protein-coding genes are not predicted to be in continuous open reading frames but are transcribed and translated into proteins

Annotation of the protein-coding genes in the *P. citri* mitogenome was done using sequence similarity searches to the *P. manihoti* mitochondrial proteins. We found a region of the *P. citri* genome with significant similarity to all 13 protein-coding genes typically found in animal mitogenomes (Fig.1A). Nine of the protein coding genes (ATP6, ATP8, COX2, COX3, CYTB, ND2, ND3, ND6, and ND4L) had continuous open reading frames (ORFs) bounded by canonical start/stop codons as found in other invertebrate mitogenomes (Supp. Fig. 1). However, the remaining four genes (COX1, ND1, ND4, and ND5) did not occur in the genome as continuous ORFs. Instead, these genes have multiple coding regions with similarity to a *P. manihoti* gene. These sequences were either found with fragments in multiple reading frames or contained premature stop codons if translated in a single frame (Supp. Fig. 1).

**Figure 1.**
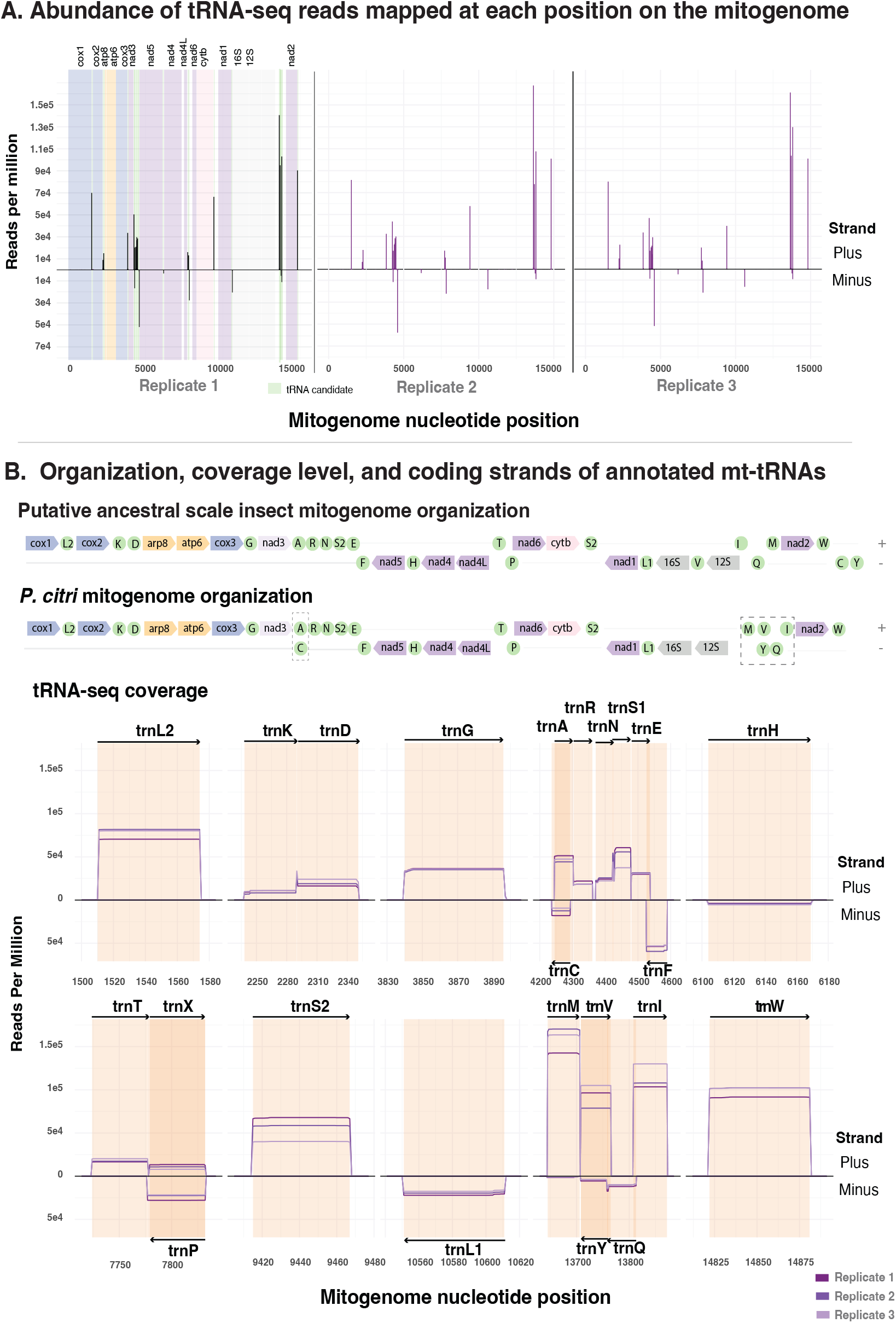
Abundance of CCA-tailed reads mapped to the *P. citri* mitogenome. **A.** YAMAT-seq read abundances globally mapped across the mitochondrial genome. YAMAT-seq will only capture RNAs with a CCA-overhang and will therefore almost exclusively sequence mature tRNA molecules. Three independent YAMAT-seq libraries were generated from purified mitochondrial fractions. The first replicate plot shows the final protein coding annotations as colored bounds (see protein annotation section for details), and candidate tRNAs are colored in green. Reads aligning to the forward strand are plotted above the x-axis (positive y-values), whereas reads aligning to the reverse strand are plotted below the x-axis (negative y-values). Bars are drawn with a width of 50 bp to visualize local read density, and the average mapped read length is 58 bp. **B**. Mitogenome organization of the ancestral scale insect and the final annotation of the *P. citri* mitogenome. Mt-tRNA rearrangements are indicated with dashed boxes. **C**. Detailed read abundance for each of the annotated mt-tRNAs. Darker orange regions indicate where two tRNAs share nucleotides (overlap).

To test whether these discontinuous gene hits were an artifact of sequencing/assembly errors or were truly reflected in the mRNAs, we performed full-length mRNA sequencing (IsoSeq) and mapped these reads to the *P. citri* mitogenome (Supp. Fig. 1). Mapped IsoSeq reads matched the reading frames of the mitogenome assembly, ruling out genome assembly errors.

ORFs generated from IsoSeq reads followed the same pattern as the mitogenome search, resulting in nine protein coding genes existing in a single frame, and four others being fragmented in multiple ORFs.

The finding of discontinuous ORFs for these protein-coding gene raises multiple possibilities. The genes could be pseudogenes, although it would be extraordinary to lose these respiration-related genes in an aerobic mitochondrion (13). Alternatively, codon reassignment, ribosomal frameshifting, transcriptional slippage, or true gene fragmentation and assembly have all been reported in mitochondria and could also be occurring in *P. citri* to restore full-length proteins/complexes. We searched the regions suspected to have frameshifts for any common codon and/or sequence patterns that could serve as translational exceptions and failed to find any exact motif at all suspected frameshifts. There was, however, a homopolymer of As or Ts of various lengths in the predicted frameshift regions for multiple genes. Both the ORFS for each gene as well as the predicted frameshift positions have been annotated in Supp. Table 1.

**Table 1.** A/T nucleotide composition across diverse A/T-biased genomes. Shown are overall A/T percentages for genomes, protein-coding sequences, and tRNA genes from mitochondrial, plastid, apicoplast, and bacterial endosymbiont genomes.

| Species | Organism | Genome | Accession | Overall A/T% | Protein coding A/T% | tRNA A/T% |
| --- | --- | --- | --- | --- | --- | --- |
| *Planococcus citri* | Insect | Mitogenome | PZ166947 | 90.9 | 88.9 | 90.6 |
| *Phenacoccus manihoti* | Insect | Mitogenome | NC_066716 | 89.3 | 88.8 | 86.1 |
| *Drosophila melanogaster* | Insect | Mitogenome | KJ947872 | 82.2 | 77.2 | 77.1 |
| *Tetranychus urticae* | Arachnid | Mitogenome | NC_010526 | 84.3 | 84.0 | 84.1 |
| *Ulva prolifera* | Green algae (Chlorophyta) | Plastid | NC_036137 | 75.2 | 74.5 | 48.0 |
| *Balanophora laxiflora* | Flowering plant | Plastid | KX784265 | 87.8 | 91.1 | 70.7* |
| *Plasmodium falciparum* | Apicomplexan | Apicoplast | LR605956 | 85.8 | 89.1 | 73.3 |
| *Shikimatogenerans sp. Tmey* | Bacterial endosymbiont (Flavobacteriales) | Endosymbiont | CP146597 | 87.9 | 88.2 | 61.3 |
| *Zinderia insecticola* | Bacterial endosymbiont (Burkholderiales) | Endosymbiont | CP002161 | 86.5 | 86.8 | 55.8 |

Finally, we performed tandem mass spectrometry (MS/MS) on purified mitochondrial proteins and detected peptides from COX2, COX3, CYTB, ND3, the first ORF of ND1, the second ORF of COX1, and the second ORF of NAD4 (Supp. Table 2). These data demonstrate that both proteins encoded on a single continuous frame and those with multiple ORFs are being translated into proteins and suggests that none of the expected 13 protein coding genes are functionally missing from the mitogenome despite some genes not being encoded in canonical or single ORFs.

### The *P. citri* mitogenome contains tRNA genes undetectable by tRNA search programs but identifiable from full-length tRNA sequencing

We scanned the *P. citri* mitogenome with tRNAscan-SE (14) which resulted in only two tRNA predictions (tRNA-Ile and tRNA-Leu(TAA)). Using the mitochondrial-specific annotation tool MITOS2 (15) resulted in 17 predicted tRNAs but with multiple predictions having poor e-value scores.

Combined, these computational approaches found no putative tRNA-Ala, tRNA-Gln, tRNA-Leu(TAG), tRNA-Tyr, or tRNA-Val (Supp. Table 3), possibly due to the extreme A/T bias of the mitogenome and/or degenerate tRNA structure of the putatively encoded tRNAs.

In order to identify tRNAs not predicted using computational approaches and find the exact genomic start/stop coordinates of each tRNA, we performed full-length tRNA sequencing using YAMAT-seq (16) on total RNA isolated from gradient-purified P. citri mitochondria (Fig. 1A). YAMAT-seq uses splint-assisted ligation with adapters that complement the CCA overhang found on all mature tRNAs (16). Read abundances from YAMAT-seq libraries can be highly biased due to inefficient reverse transcription of some tRNAs due to base modifications (17). However, because both adaptors are ligated directly to mature tRNAs during library preparation, full-length tRNA sequences are amplified during sequencing library creation and can be used to determine tRNA gene bounds.

YAMAT-seq libraries from purified *P. citri* mitochondria resulted in 23 clear ‘hotspots’ in the mitogenome corresponding to region with a large percentage of mapped reads (Fig. 1A). From these regions with abundant YAMAT reads, 23 candidate tRNA sequences were identified, and the most frequent first and last positions for each sequencing peak were assigned as the gene’s boundaries.

All 17 of the MITOS2 predicted tRNAs corresponded to a hotspot region with high read abundance (Supp. Table 3). For 16/17 of those tRNAs, the most abundant start/stop read coordinates were within 4 bp of the computationally predicted start/stop coordinates (Supp. Table 3). Only tRNA-Arg was substantially different than the predicted sequence at 10 bp shorter. Although tRNA-Gln, tRNA-Leu(TAG), and tRNA-Tyr were not identified in the mitogenome when using computational approaches, we found a highly expressed region of the mitogenome that contained the corresponding anticodon to each of these missing tRNAs (Fig.1B, Supp. Table 3). These sequences were BLASTed to mt-tRNA databases from close relatives (Supp. Table 3) to test for sequence similarity to other mt-tRNAs. By taking a combination of the best hit in conjunction with RNA folding strategies (see section on tRNA structure below) the tRNA-Gln, tRNA-Leu(TAG) and tRNA-Tyr were identified and annotated (Fig. 1B).

### The mitogenome encodes multiple tRNA ‘mirrors’, tRNAs produced from opposite strands of a single genomic location

Surprisingly, three genomic locations with a putative tRNA gene had substantial YAMAT expression from both strands, and both orientations were post-transcriptionally modified with a CCA tail (Fig. 2, Supp. Table 3). These mirrored tRNAs carry reverse-complement anticodons: tRNA-Cys(GCA) mirrors tRNA-Ala(TGC), and tRNA-Tyr(GTA) mirrors tRNA-Val(TAC) (Fig. 3) Two of these mirrored tRNAs, tRNA-Ala(TGC) and tRNA-Val(TAC), contain the two anticodons missing from the rest of the predicted tRNA set. As the expected 22 mt-tRNAs can only be accounted for by including the bidirectional expression of these two tRNA loci (tRNA-Tyr/Val and tRNA-Cys/Ala), these data raise the possibility that a single tRNA locus is being transcribed to produce two distinct tRNAs, one from each strand. The tRNA-Pro(TGG) locus also had substantial antisense expression (designated as trnX, Fig. 2), but a tRNA with the anticodon CCA (decoding tryptophan) is not typically found in animal mitochondria (18), making the function of this transcript unknown. All other genomic locations with a computationally or experimentally predicted tRNA gene only had appreciable sequence coverage arising from one strand of the genome (Fig. 2).

**Figure 2.**
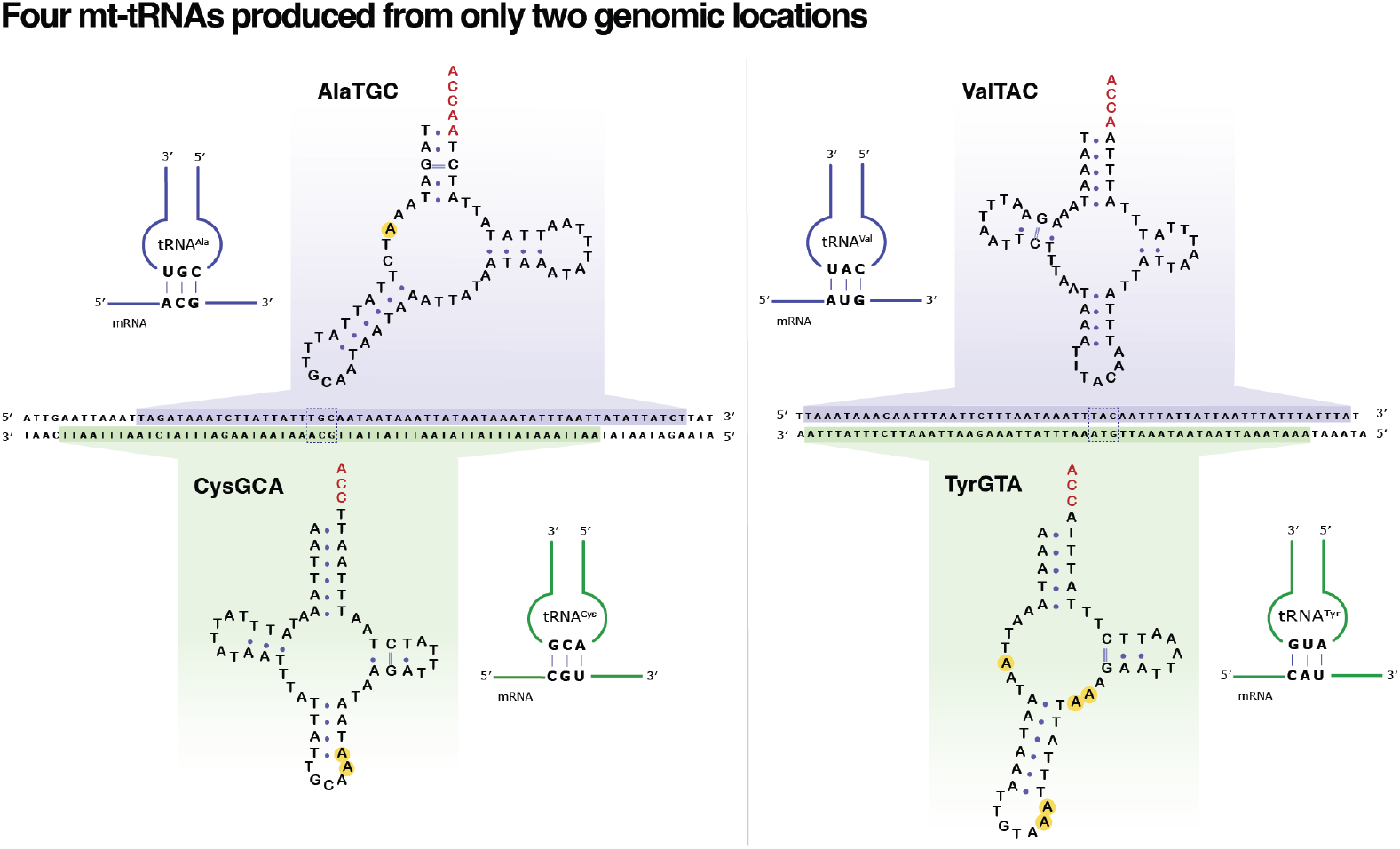
Two loci in the mitogenome produce four tRNAs with different anticodons. The genomic sequence and transcribed strands for each of the mirror mt-tRNAs, Cys/Ala and Tyr/Val. Structures were manually folded to maintain stem and loop features and presented above or below the genomic locus in the mitogenome. Post-transcriptionally added nucleotides are indicated in red. Nucleotides with a yellow circle have evidence of a base modification that interferes with reserve transcription. The genomic coordinates of the anticodon nucleotides (indicated with a dashed box) are the same for each pair of bidirectional tRNAs but are reverse complements of each other and therefore decode for different amino acids. Folded structure diagrams were created using VARNA ver. 3.9 (67).

**Figure 3.**
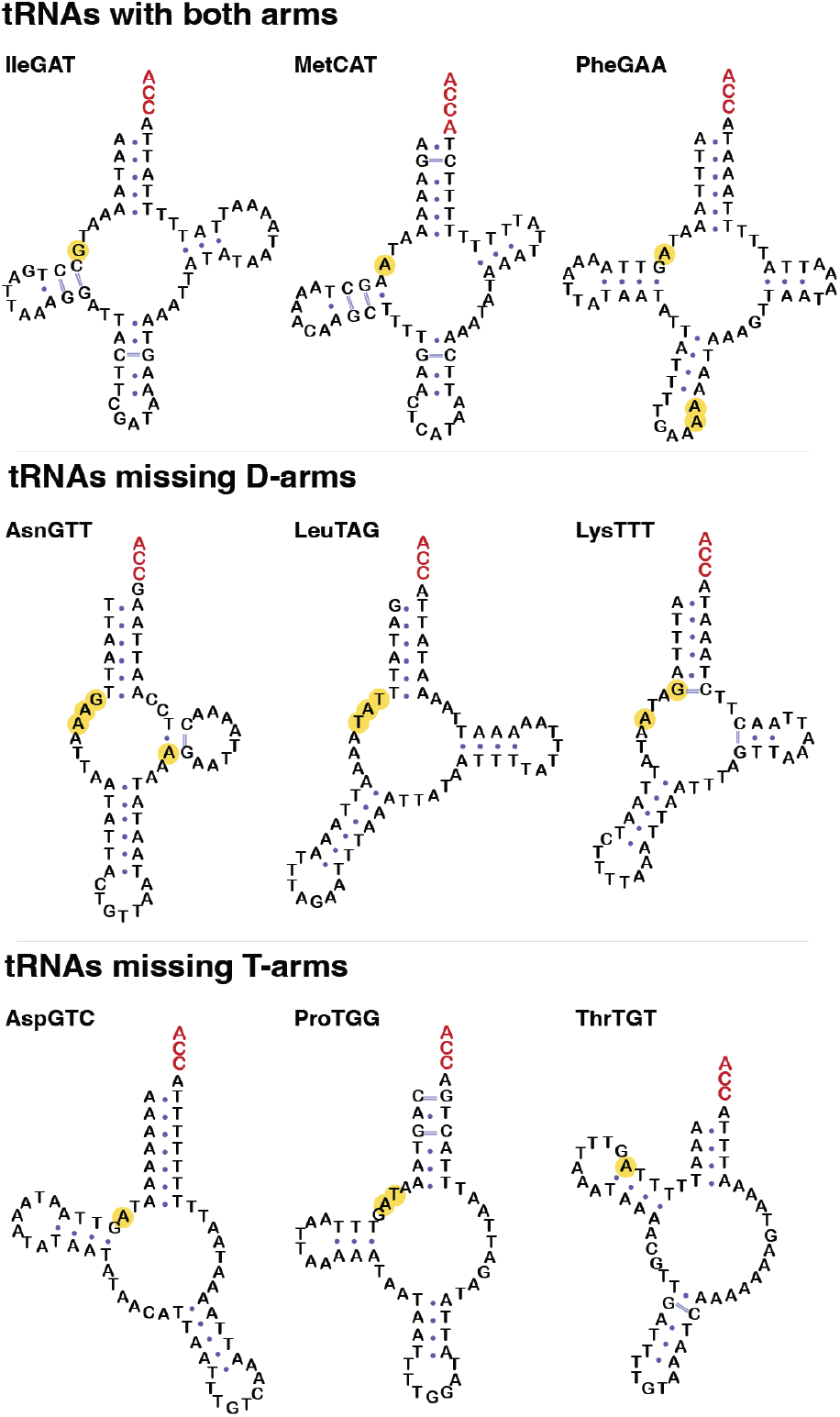
Predicted folding of nine representative mt-tRNAs. Post transcriptionally added nucleotides are indicated in red. Nucleotides with a yellow circle have evidence of a base modification. RNA folding was done manually to maintain stem structures and visualized using VARNA ver. 3.9 (67).

Although very rare in bilaterian animals, mt-tRNA gene loss and functional replacement through the import of nuclear-encoded tRNAs does occur in many other eukaryotic lineages (5, 19). To test for the possibility that *P. citri* has functionally lost mt-tRNAs and is now importing nuclear-encoded tRNAs, we performed differential abundance analysis on gradient-purified mitochondrial samples vs. total insect samples to assess if any tRNAs not encoded on the mitogenome were enriched in *P. citri* mitochondria. A database of all *P. citri* nuclear, mitochondrial, and bacterial endosymbiont tRNAs (*P. citri* has two nutritional endosymbionts encoding their own tRNAs) was assembled (Supp. Table 4), and YAMAT libraries were sequenced from three total insect and gradient-purified mitochondria samples and mapped to this database (all mapped counts for all YAMAT-seq libraries in Supp. Table 5). As expected, all the mitochondrial tRNAs were highly enriched (>19 fold) in mitochondrial fractions; however, no nuclear tRNAs approached that enrichment level in the analysis (Supp. Fig. 2, Supp. Table 6). The failure to identify strong candidates for preferential import of any nuclear-encoded tRNAs suggests that *P. citri* mitochondria still rely on natively encoded mt-tRNAs for translation despite the extremely degenerate nature of the mt-tRNAs.

### *P. citri* mt-tRNAs are the most A/T biased tRNAs reported

With an average length of only 58 bp, the majority (15/22) of the mt-tRNAs are too short to produce a canonical cloverleaf tRNA shape with both D- and T-arms (Fig. 3, see Supp. Fig. 3 for all mt-tRNA models). The truncated and extreme A/T bias of these tRNAs prevents RNA folding algorithms such as RNAfold (20) from predicting the expected stem and loop structures found in tRNAs; therefore, we manually folded all tRNA sequences to maintain these features (Fig. 3 and Supp. Fig. 3). Although many of the tRNAs are represented as missing only one arm (retaining either a D- or T-arm), these stems were often short (> 4 bp) and sometimes contained modified bases that would interfere with base pairing (see section on base modification below), raising the possibility that some of these tRNAs may actually be armless in vivo. The A/T composition of the tRNAs is extraordinarily biased, with a range of 82-96% A/T and an average of 90.6% A/T (Table 1, Supp. Table 3). Remarkably, tRNA-Leu(TAG) is almost 97% A/Ts, having only 1 G and 1 C that are not part of the non-templated CCA tail, and lacks even a single C-G base pair (Fig. 3).

### Mirror mt-tRNAs expressed from opposite strands have unique modification and editing profiles

All tRNAs typically have multiple modified nucleotides (21, 22), and these modifications can play critical roles in tRNA structure and function. Because some base modifications can inhibit reverse transcriptase (RT) activity, modifications can be mapped to specific nucleotides in a tRNA through the detection of RNA polymerase errors (nucleotide misincorporations and indels) in the resulting cDNA (23–25). These misincorporation signatures can then be used to infer both the position and the identity of the modification, producing a modification map or ‘index’ of a tRNA. We applied a modification detection pipeline to the *P. citri* mt-tRNAs to produce a tRNA modification index by globally aligning mapped reads to each reference using MAFFT (26) and calculating percentage of mismatched reads for each position along the tRNA (see Supp. Table 7 for all aligned base calls).

The number of detected modifications for the mt-tRNAs was low, with an average of only 1.87 detected base modifications per tRNA when a 20% cDNA misincorporation threshold was applied. There was, however, a consistent signal of a base modification in 5’ region of the mt-tRNAs (Fig. 4, Supp. Fig. 4). All but one mt-tRNA had evidence of one or multiple A or G modifications within positions at 7-9 of the tRNA (Supp. Fig. 4). One likely candidate is 1-methyladenosine (m1A) which is a very common base modification in mt-tRNAs and is known to cause RT-inhibition signatures (27). Only three mt-tRNAs (Cys, Phe, and Tyr) had RT-inhibiting base modifications in the anticodon loop, and in all cases, the misincorporation signal was a two base pair deletion immediately following the anticodon. The modifications N6-isopentenyladenosine (i6A), or the modified i6A, 2-methylthio-N6-isopentenyladenosine (ms2i6A), are known to occur on animal Cys, Phe, and Tyr mt-tRNAs at this position (27), and their bulky nature often cause reverse transcriptase stalling/skipping instead of misincorporating the wrong nucleotide (28).

**Figure 4.**
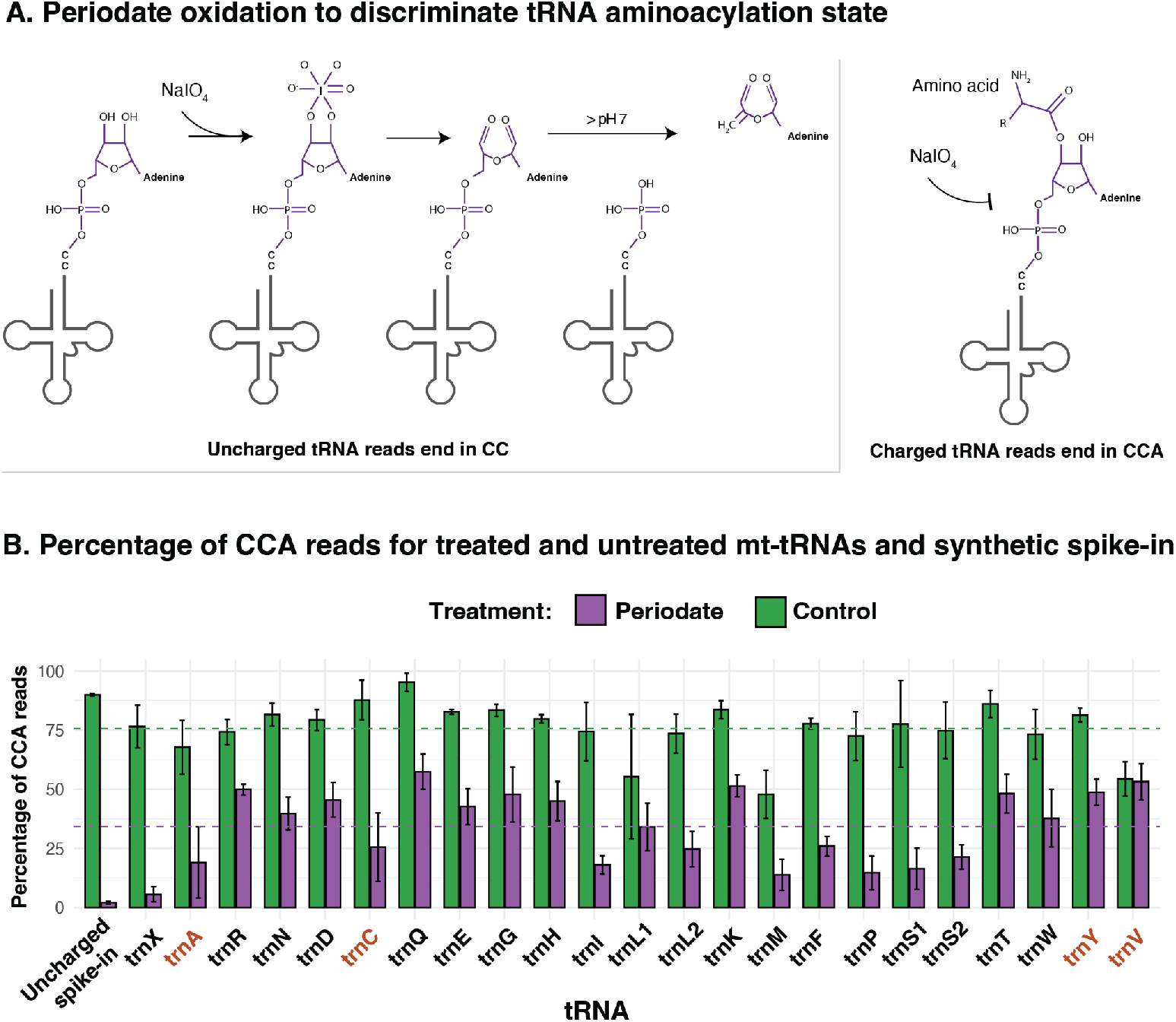
Using periodate oxidation to determine the proportion of charged mt-tRNAs in *P. citri* mitochondria. **A.** Graphic summary illustrating the use of the Whitfeld reaction followed by 3′ adenosine cleavage to discriminate between deacylated and acylated tRNAs. For uncharged tRNAs the 3′ adenosine is oxidized by periodate and then cleaved off by borax-induced β-elimination. Cleavage of the terminal adenosine means that resulting tRNA-seq reads from uncharged tRNAs end in CC. For charged tRNAs, the 3′ adenosine protected from periodate oxidation and reads will end in CCA. **B**. Percentage of reads with a complete CCA tail (CCA/(CC+CCA reads) for each mt-tRNA reference and an uncharged, synthetic tRNA. Three total RNA samples from gradient-purified P. citri mitochondria were split into periodate and untreated subsamples. Bars represent the average percentage of CCA reads across the three replicates. Error bars indicate ± 1 SD across replicate libraries. Dotted green line is the mean percentage (75.7%) of CCA reads in untreated (control) libraries. Dotted purple line is the mean percentage (34.2%) in periodate treated libraries. Mt-tRNAs in red are the mirror pairs.

Notably, the two pairs of mirrored tRNAs (Ala/Cys, and Tyr/Val) have very different predicted secondary structures and entirely different modification profiles from their complementary sequence (Fig. 2). Mt-tRNA-Ala has evidence of a base modification at position 8A, whereas its tRNA-Cys mirror lacks a modification at any position in the 5’ region, and instead has the bulky modification immediately following the anticodon mentioned previously. Mt-tRNA-Tyr is the most modified mt-tRNA in these data, with five different modifications (Fig. 2, Supp. Fig. 3) whereas its mirror tRNA-Val lacked a strong signal for base modification at any position (Fig. 2).

In addition to base modifications, mt-tRNAs can also have bases added post-transcriptionally to produce a final sequence that is different than the sequence in the mitogenome. One example that has been reported in multiple animals is the addition of adenosine at the 3′-end of mt-tRNAs prior to CCA tailing (29). This type of editing often occurs when partially overlapping tRNA genes produce truncated sequences that are then post-transcriptionally “finished” to restore a full-length tRNA. In *P. citri*, mt-tRNA-Gln, Gly, Met, Thr, and Val all had the addition of a single adenosine at the 3′-end, and the tRNA-Ala had the addition of two adenosines (Supp. Table 2).

### Mirror mt-tRNAs expressed from opposite strands are aminoacylated

For a tRNA to be functional in translation it must first be aminoacylated, or “charged,” with an amino acid to the terminal adenosine in the CCA tail. Charged tRNAs can be quantified using recently developed tRNA sequencing methods that use the Whitfeld reaction to remove the terminal 3′ nucleotide of uncharged tRNAs that lack the protective amino acid (30, 31). The percentage of reads ending in a CC (not aminoacylated) vs those ending in the complete CCA (aminoacylated) can then be used to measure the proportion of aminoacylated molecules for a specific tRNA (Fig. 4A). We used this method to determine which of the identified sequences in the P. citri mitochondria could function in translation and test whether both mirror tRNAs produced from a single locus were aminoacylated.

Three biological replicates from gradient-purified *P. citri* mitochondria were used to generate and sequence multiplex small RNA-seq (MSR-seq) libraries (30). Each replicate was split into two subsamples where one subsample was treated with periodate to distinguish between aminoacylated and uncharged tRNAs, and the other served as a negative control with no periodate treatment. This negative control provides insight into the base level of CCA tailing and any CCA tail damage during extraction. The average percentage of CCA reads in these untreated libraries was approximately 74% for the mt-tRNAs (Fig. 4B, Supp. Table 8). A synthetic Bacillus tRNA-Ile(GAT) served as an uncharged tRNA control and was added to all samples after RNA extraction, but before the beginning of library preparations to measure the effectiveness of periodate treatment in removing the terminal A nucleotide.

One difficulty in applying these aminoacylation detection methods to the P. citri mt-tRNAs was the extreme A/T bias and the necessary number of PCR cycles used to amplify MSR-seq libraries. PCR and Illumina sequencing have known biases against A/T rich sequences, resulting in lowered or even elimination of A/T biased sequences in the final read population of complex libraries (32). When applying 15 cycles of PCR, periodate treated libraries had pronounced drop out of the most A/T rich sequences and those mt-tRNAs expected to have the lowest expression (Supp. Fig. 5). However, when lowering the number of PCR cycles to 10 or 6 cycles and using a barcoding and library pooling scheme, we were able to detect all 23 mt-tRNA references in treated and untreated libraries (Supp. Fig. 5). Periodate treatment was very effective at removing the terminal A in the uncharged tRNA spike-in control, with an average of only 2% of the reads ending in the complete CCA in the treated libraries (Fig. 4B). In contrast, the average percentage of mt-tRNA reads ending in CCA was 34.2% (Fig. 4B, Supp. Table 8B). Across all mt-tRNAs, periodate treatment revealed a strong and highly significant protection relative to the uncharged Bacillus spike-in (FDR<< 0.05, binomial glm), with odds ratios ranging from 21-fold (tRNA-Cys) to greater than 400-fold (tRNA-Val) (Supp. Table 8C). Mt-tRNA-Cys and mt-tRNA-Ala had the fewest reads mapped, resulting in high standard deviation of the read classes. However, the average percentage of reads with the complete CCA tail for both mt-tRNA-Cys and mt-tRNA-Ala was still much higher than the negative control with 25.5% and 19.0%, respectively (Fig. 4). The detection of mt-tRNA-Tyr and mt-tRNA-Val sequences was marginally better and had higher-than-average CCA read percentages with 48.7% and 53.2%, respectively. The antisense expression of tRNA-Pro (trnX) was an interesting case, because although this sequence was statistically more protected than the uncharged spike-in (Supp. Table 8C), it still showed very little retention of the complete CCA tail, with an average of 5.6% of the reads containing the terminal A across the replicate libraries. This low rate of protection provides evidence that this RNA is not significantly aminoacylated and may not be participating extensively in translation as a true tRNA.

**Figure 5.**
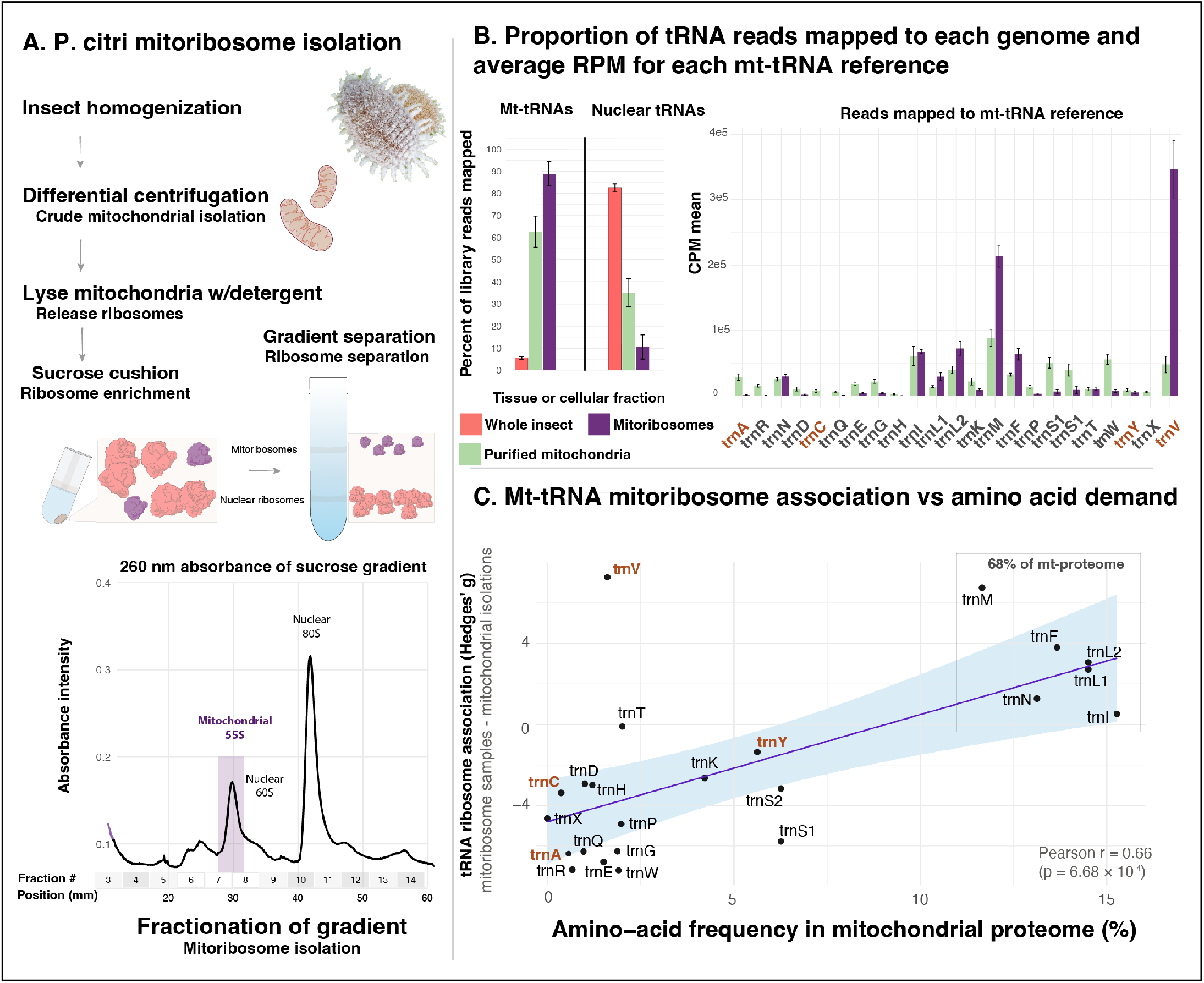
Mitoribosome tRNA sequencing to assess mt-tRNA occupancy. **A.** Whole mealybugs at multiple life stages were homogenized and subjected to differential centrifugation to enrich for mitochondria. The mitochondrial fraction was lysed with detergents to release mitoribosomes, which were then separated by a sucrose cushion and continuous sucrose gradient to isolate mitoribosomes. The bottom panel shows a representative 260 nm absorbance trace from a sucrose gradient, with the mitochondrial monosomes (intact ribosomes) highlighted in a red box; nuclear large subunit (60S) and monosome (80s) peaks are labeled in text. Total RNA was extracted from the mitoribosome-containing fractions (fractions 6–8) and used to construct mitoribosome tRNA-seq libraries. **B**. Left plot: mean percentage of tRNA-seq reads mapping to the nuclear and mitochondrial genomes from whole-insect, gradient-purified mitochondrial and mitoribosome libraries. Right plot: normalized counts per million (CPM) reads mapping to each mt-tRNA reference from purified mitochondrial and mitoribosome libraries. Error bars indicate ± 1 SD across replicate libraries. Mirror tRNAs are indicated with red text. **C**. Each mt-tRNA’s ribosome association versus the frequency of its cognate amino acid in the mitochondrial proteome. Ribosome association was calculated as Hedges’ g (mean-difference statistic) for each tRNA, comparing abundance (normalized CPM reads) in mitoribosome libraries versus total mitochondrial libraries. Positive g values indicate higher relative abundance of a tRNA in the ribosome-associated fraction; negative values indicate enrichment in the total mitochondrial libraries. The x-axis shows the frequency of the corresponding amino acid in the mitochondrial proteome. Association strength was evaluated by simple linear regression (g ~ amino-acid frequency). Each point represents a single mt-tRNA; the solid line shows the fitted regression with its 95% confidence interval (light-green shading). The two pairs of tRNAs with bidirectional expression are highlighted in orange. The boxed region indicates that 68% of the amino acids in the coding sequences are composed of only five amino acids. Amino acid frequencies were determined using homology searches and full-length mRNA sequencing (see protein-coding section below). Mealybug photo in A by Alexander Wild.

We also calculated the percentage of reads ending in CCA for nuclear tRNAs in these libraries (Supp. Fig. 6) and found an overall average of 37%. The average percentage of reads ending in CCA for all P. citri tRNA references can be found in Supp. Table 8. Overall, these data support the hypothesis that *P. citri* maintains 22 aminoacylated mt-tRNAs.

### Mirror mt-tRNAs are found in mitochondrial ribosome isolates, with mt-tRNA-Val found in very high abundance

To test the occupancy of mt-tRNAs in the mitochondrial ribosome, we performed mitoribosome tRNA-seq by isolating mitochondrial ribosomes through sucrose density gradient centrifugation followed by YAMAT-seq on RNA isolated from those fractions. While raw abundance values from tRNA sequencing methods likely do not reflect absolute biological abundances due to reverse transcription stalling from tRNA modifications, relative comparisons across samples and treatments using the same tRNA-seq method can provide meaningful abundance differences. Using this logic, we next measured whether the tRNA population in whole mitochondria was different than the tRNA population purified with mitoribosomes, if these differences reflected amino acid demand, and if the mirror tRNAs were abundant in mitochondrial ribosome fractions.

Mitoribosomes are substantially smaller than cytosolic ribosomes and can be separated based on differences in sedimentation (33). We separated *P. citri* mitoribosomes from cytosolic ribosomes using differential centrifugation followed by sucrose gradient fractionation (Fig. 5A). Gradients were fractionated while monitoring absorbance at 260 nm to identify ribosome-containing fractions. The absorbance profile showed multiple peaks, including a distinct peak preceding the large cytosolic ribosome peak (Fig. 5A). To verify ribosome separation, total RNA was extracted from each fraction and RT– qPCR was performed targeting the large rRNA from each ribosome class (primers can be found in Supp. Table 9). Mitochondrial rRNA was most abundant in fractions 6–7 (Supp. Fig. 7). RNA from fractions 6–8 was therefore pooled for mitoribosome tRNA sequencing. With an average of 88.9% of the reads from the mitoribosomes libraries mapping to a mt-tRNA reference, these mitoribosome tRNA-seq libraries were highly enriched for mt-tRNAs compared to whole insect, or even when compared to gradient-purified mitochondria libraries (Fig. 5B). The relative abundance of each mt-tRNA changed considerably in isolated (total) mitochondria libraries compared to mitoribosome libraries. Most notably, there was an extreme abundance of reads mapping to mt-tRNA-Val in mitoribosome samples, representing an average of 38.8% of the reads (Fig. 5B). As animal mitochondrial ribosomes have been reported to use mt-tRNA-Val as a structural component of the ribosome itself (replacing the 5S rRNA) (34), this high abundance of mt-tRNA-Val may reflect its role as a structural component of the mealybug mitoribosome.

To test whether mitoribosome tRNA-seq abundances reflect amino acid usage in the mitochondrial proteome, we computed Hedges’ g (a small-sample size corrected Cohen’s d) for each tRNA using CPM-normalized read counts and comparing mitoribosome to total mitochondrial read abundances. We then regressed this mean difference statistic (Hedges’ g) on the frequency of the cognate amino acid in the mitochondrial proteome (Fig. 5C). This was done under the premise that the standing tRNA population in whole mitochondria would be different than the population loaded in the mitoribosomes, and that this difference should reflect amino acid demand. Association strength was evaluated by simple linear regression (g ~ amino-acid frequency) and found to have a Pearson r correlation of 0.65 (p = 0.00108) and Spearman correlation coefficient of 0.63 (p = 0.00133), representing a moderately strong relationship between tRNA-seq abundance in the mitoribosome samples and amino acid use. The mitochondrial proteome in P. citri is extremely biased as a consequence of the extreme A/T composition of the genome. Only five amino acids (Ile, Leu, Phe, Asn, and Met) make up over 68% of the amino acids in the proteins encoded on the mitogenome, and the tRNAs that carry these amino acids had a much greater change of abundance in the mitoribosome samples than the other tRNAs (Fig. 5C). 13/22 mt-tRNAs were within a 95% confidence interval (CI) of expected mitoribosome occupancy (Fig. 5C). The amino acids Ala and Cys are the least common amino acids in the mitochondrial proteome (representing 0.57% and 0.37% respectively), but both tRNAs originating from the tRNA-Cys/Ala locus were found to be in the CI of expected abundance change in mitoribosome samples. In contrast to the extreme overrepresentation of mt-tRNA-Val, the mirrored sequence at this locus, tRNA-Tyr, was solidly in the CI of expected abundance change in mitoribosome samples (Fig. 5C).

## Discussion

The complementary expression of two tRNAs from a single locus is a major departure from the known tRNA gene organization of all types of genomes and organisms. There are rare examples of two proteins being encoded with bidirectional expression of a locus producing two different proteins (35, 36), but the majority of functional antisense gene expression is the generation of a noncoding RNA complementary to mRNA with modulating effects on its translation (37, 38). The complementary nature of DNA/RNA double strands and the stem-loop, palindromic structure of tRNAs may make them amenable to this unusual form of mirrored expression.

Antisense expression is ubiquitous in animal mitogenomes, as both strands are fully transcribed and later processed by RNA maturation enzymes to release individual RNAs from the polycistronic precursor (39). Typically, these transcriptional byproducts are rapidly degraded and maintained a very low level (40, 41). Nevertheless, the origin of mirror tRNAs could evolve in a straightforward manner, where a spurious, antisense transcript of a tRNA has enough structural similarities to be recognized by tRNA maturation enzymes and processed like a bona fide tRNA, thereby avoiding the degradation pathway. For example, the CCA-adding enzyme has been shown to act on non-tRNA substrates that have a stem-loop structure (42, 43). If these processed sequences can then be recognized by an aaRS, the result would be a pool of aminoacylated mirror tRNAs alongside the ancestral mt-tRNA. The participation of these mirror tRNAs in ribosomal decoding would effectively make the ancestral tRNA gene redundant, eventually allowing for its eventual loss and making the system functionally dependent on the bidirectional expression of one tRNA locus.

The presence of small (1-3 bp) overlaps in mt-tRNA genes is common in animal mitogenomes (44). These overlaps often result from truncated acceptor stems which are then repaired through the post-transcriptional addition of 3’ nucleotides (29, 45–47). An extreme situation of mt-tRNA gene overlap has been suggested for the isopod *Armadillidium vulgare* where the mt-RNA-Cys and mt-tRNA-Tyr genes are predicted to overlap by 20 bp on the same strand, and the mt-tRNA-His and mt-tRNA-Phe genes are predicted to overlap by 19 bp on the opposite strand (48). The sequencing of RT-PCR products from the mt-tRNA-His confirmed the presence of a CCA tail in addition to extensive 3’ editing of nucleotide additions not encoded in the mitogenome (48). Importantly, though, no full-length tRNA sequencing has been performed to confirm the coordinates of the potentially overlapping genes.

Regardless of whether mt-tRNA gene overlap is frequently occurring at only a few base pairs followed by repair or at a larger scale, it is difficult to imagine a scenario whereby tRNA mirrored expression could evolve through a “creeping overlap” mechanism. Notably, although some *P. citri* mt-tRNAs are being edited though the addition of 1-2 non-templated As (Supp. Table 3), the mirror tRNAs we report here do not arise from truncated transcripts which are then edited to full-length tRNAs: they are encoded on the genome as overlapping tRNA genes which share completely overlapping anticodons (Fig. 2).

We suspect that the centering of both tRNAs around the same genomic location would have been nearly impossible through a progressive shift of transcription. Instead, the wholesale bidirectional expression of a single locus followed by slightly different cleavage points from the polycistronic transcript would give rise to two tRNAs that have mirrored anticodons.

Determining the identity of the ancestral (and lost) mt-tRNA genes in *P. citri* was complicated by the truncated and incredibly A/T biased nature of the sequences. This difficulty was particularly true for the mt-tRNA-Cys/Ala locus. For consistency, we used the MITOS2 call for the identification of the ancestral tRNA at this mirror position, but the score was quite low (Supp. Table 2). There is, however, evidence from gene synteny that this locus was ancestrally tRNA-Ala in scale insects (Fig.1B). Additionally, insects have a conserved 3G-C70 base pair in mt-tRNA-Ala (48), which is present in the P. citri sequence (position 3G-57C, Supp. Fig. 4) and this G-C base pair would not exist in antisense expression of the ancestral mt-tRNA-Cys. For tRNA-Val/Tyr mirrors, there was not strong syntenic evidence for the identity of the tRNA at this position (Fig.1B), but this sequence showed significant similarity to previously annotated tRNA-Tyr in multiple scale relatives (Supp. Table 3).

The high abundance of this tRNA in the mitoribosomes fractions suggests that mt-tRNA-Val is a structural component of the mitoribosomes like in mammals (50). If true, that would mean that the mirrored expression of the mt-tRNA-Tyr gene has replaced both the mt-tRNA-Val in decoding as well as a structural component of the mitoribosome.

The widespread antisense/mirror expression and CCA-tailing of mt-tRNA genes was first reported in the unrelated spider mite *Tetranychus urticae*, where nine mt-tRNAs were found to have substantial anti-sense expression and posttranscriptional modification (12). However, the mitogenome of *T. urticae* still encodes the expected 22 mt-tRNA genes, making the complementary expression of mt-tRNAs redundant (all the anticodons are still encoded by a separate tRNA gene) and thus the functional significance of these mirrors unknown. It would be interesting to determine if these anti-sense tRNAs are aminoacylated in *T. urticae* as this species may represent an intermediate evolutionary step prior to the loss of one of the mt-tRNA genes as reported in *P. citri*.

Notably, *T. urticae* mt-tRNAs are also truncated and A/T rich like that of *P. citri*. In fact, multiple animal lineages have independently evolved short mt-tRNAs structures (44). In addition to multiple arachnid lineages of spiders and mites (11, 51–53), a class of nematodes (Enoplea) has also been reported to have highly degenerate mt-tRNA structures (54, 55). The mitogenome of the nematode *Romanomermis iyengari* has some of the shortest tRNAs reported and is actually annotated as having mirror Ala/Cys mt-tRNAs (10). However, subsequent computational searches in Enoplea mitogenomes found 22 separate mt-tRNA candidate genes without needing to postulate the existence of mirrors (10). Difficulty in predicting degenerate mt-tRNAs has resulted in numerous annotations missing mt-tRNA genes below the expected 22 and highlights the importance of generating experimental evidence for these essential gene products. But the functional expression of mirrored mt-tRNAs in *P. citri* and the presence of antisense/mirror tRNA sequences in the unrelated mite *T. urticae* raises the fascinating possibility that tRNA mirroring is an evolutionary route for mt-tRNA gene loss and may have convergently evolved multiple times independently.

The theoretical existence of mirror mt-tRNAs has been proposed both in extant systems (56) and in the ancestral decoding apparatus of early life (57, 58).The Rodin and Ohno hypothesis proposes that the two major classes of aminoacyl-tRNA synthetases (Class I and II) originated from complementary strands of a common ancestral gene, and that these enzymes charged mirror tRNA molecules. Rodin and Ohno highlight the striking structural symmetry that Class I and Class II aaRSs interact with opposite grooves of the tRNA acceptor stem, and the codons they recognize correspond to complementary tRNA anticodon triplets, dividing the genetic code along a strand-symmetric axis (57). The discovery that mirror tRNAs have evolved in a bilaterian animal mitogenome that experiences strong genetic drift and a high substitution rate may offer insight into the limits of a minimal translation system and a window into the ancient and ancestral core translation machinery. As drift becomes a greater force in the evolution of genomes, only the most deleterious mutations are selected against (59). This process may eventually lead to mutational meltdown (60) but may also result in the winnowing of a system down to its most essential components (61). Therefore, the reduction in the number of tRNA loci and the bidirectional expression of mt-tRNA in *P. citri* may represent an extreme form of system reduction akin to the ancestral decoding system. Evidence for reductive processes being used as windows into ancestral machinery may also be found in the reduced mt-tRNA structures themselves. The loss of both arms in mt-tRNAs in multiple lineages (including possibly P.citri) shares resemblance to the minihelix, a simple stem-loop structure proposed to be the ancestral tRNA molecule (62, 63).

Finally, the mirrored expression of tRNAs poses a striking evolutionary problem for *P. citri*: how do tRNA-interacting enzymes adapt to substrates transcribed from both strands? Because tRNAs and aaRSs are typically tightly co-evolved to ensure the faithful decoding of the genome, the existence of mirror tRNAs implies that a single genomic locus must maintain compatibility with two distinct aaRSs acting on complementary substrates. Genes predicted to encode separate mitochondrial Cys, Ala, Tyr, and Val aaRSs are present in the *P. citri* nuclear genome (Supp. Table 10), but it remains unclear how these enzymes accommodate the mirrored tRNA structures, or whether distinct structural adaptations have evolved to recognize them.

## Methods

### Crude isolation of *P. citri* mitochondria

Mealybugs were reared on small potatoes in a chamber set to 12 hr light/12 hr dark at 26°C with 50% relative humidity. Detailed *P. citri* mitochondrial isolation methods can be found in Supplementary Methods. Briefly, 2 g (~10,000 individuals) of mealybugs at all life stages were used for each mitochondrial isolation. Ethanol-cleaned mealybugs were homogenized in isolation buffer (50 mM HEPES-KOH pH 7.5, 10 mM KCl, 1.5 mM MgOAc, 70 mM sucrose, 210 mM mannitol, 1 mM EDTA, 1 mM EGTA, 1 mM DTT) using a glass dounce homogenizer. Homogenate was filtered over cheese cloth and Miracloth. Filtered homogenate was spun at 1,000 g for 5 min at 4 °C with a fixed angle rotor. Pellet was discarded and supernatant was spun again at 1,000g for 5 min at 4 °C. Pellet was again discarded, and the supernatant was filtered over 100 µm mesh filters. Cleaned supernatant was spun at 15,000g for 15 min at 4 °C to pellet mitochondria. The supernatant was discarded, and the crude mitochondria pellet was resuspended into 1 mL SEM buffer (250mM sucrose, 20mM HEPES-KOH pH7.5, 1mM EDTA, 1mM EGTA) with a paint brush. This crude mitochondrial isolate went directly into sucrose gradient purification (see below), or the pellet was snap frozen in liquid nitrogen for use in mitochondrial ribosome isolations or DNA extractions.

### Gradient purification *P. citri* mitochondria

Resuspended crude mitochondrial pellets were carefully layered on top of a 15-23-32-60% sucrose gradient (in 50mM HEPES-KOH pH7.5, 1 mM EDTA, 1mM EGTA). The gradient was spun at 28,000g for 1 hr at 4 °C in a swinging bucket rotor. Purified mitochondria formed a band at the 32/60% interface and was collected by aspiration. Purified mitochondria were washed by adding 5X volumes of SEM buffer and spun at 15,000g for 15 min. Supernatant was discarded and the pellet was snap frozen in liquid nitrogen for further analyses.

### *P. citri* mitogenome sequencing and assembly

#### Nanopore sequencing

High molecular weight DNA was extracted from a crude mitochondrial isolation and sequenced with Oxford Nanopore, using the Ligation Sequencing Kit (SQK-NBD114.24) with the NEBNext Companion Module (E7180L) to manufacturer’s specifications. Guppy1 (v6.5.7) was used for basecalling (SUP), demultiplexing, and adapter removal. Reads were trimmed with Porechop (0.2.3-seqan2.1, (--discard_middle).

#### Illumina sequencing

DNA was extracted from a crude mitochondrial isolation and sequenced with the NovaSeq X Plus sequencer with 2×151bp paired-end reads. Illumina sequencing libraries were prepared using the tagmentation-based and PCR-based Illumina DNA Prep kit. PCR amplification was done with only 5 cycles of PCR. Library was sequenced on Demultiplexing, quality control was performed with bcl-convert1 (v4.2.4). Reads were trimmed with cutadapt (-q 20 --minimum-length 50 -e 0.15 -j 24).

#### Assembly

Trimmed Nanopore reads were BLASTed (blastn -dust no -evalue 1e-7) to the previously published P. citri mitogenome (OX465514). Reads with homology (a successful BLAST hit) to the mitogenome were used in a Flye (2.8.1-b1676) assembly with 5 iterations. This assembly was then polished with Illumina reads using Perbase (https://github.com/sstadick/perbase, base-depth --max-depth 1000000), and a custom perl script (https://github.com/warrenjessica/Mealybug_mirror_mt-tRNAs/tree/main/Mitogenome_assembly). Bowtie2 v2.5.4 (64) was used to map Illumina reads to the Nanopore assembly. Assembly was deposited on GenBank: PZ166947

### Mitogenome annotation

tRNA annotation: Mt-tRNA annotation was performed using tRNA search programs and validated with full-length tRNA sequencing. The completed mitogenome assembly was scanned with tRNAscan-SE using the organellar search option (-O), and with MITOS2 under default parameters, using the Metazoa reference database and the invertebrate mitochondrial genetic code. The most frequent start and stop coordinates for each high-abundance YAMAT read were assigned as the tRNA boundaries. These reads were compared to a database of mt-tRNAs from closely related species (Supp.Table 3) using BLASTn (-task blastn, -dust no). Any tRNAs not identified by MITOS2 were assigned to the identity of their closest BLAST match from a relative.

tRNA folding: Each mt-tRNA was manually folded by positioning the predicted anticodon approximately in the center of the tRNA, and any G/C base pairs were predicted to be stabilizing stems.

Protein annotation: Protein-coding gene annotation was performed using a reference amino acid database of mitochondrial proteins from *P. manihoti* (NC_066716). This database was queried against the *P. citri* mitogenome assembly using tBLASTn (-evalue 0.1, -db_gencode 5, -seg no). Regions showing homology t *P. manihot* genes were searched for the longest ORFs. Homologous regions corresponding to all protein-coding genes were recovered except for ATP8 and ND4L. For these two genes, ORFs were identified within the syntenic regions where they occur in other scale insects, and the resulting ORFs were confirmed by BLASTp against a database of ATP8 and ND4L sequences from related species. When a gene corresponded to multiple ORFs within the *P. citri* mitogenome, each ORF was annotated as part of the gene (Supp. Table 1).

### Full-length tRNA sequencing

Full-length tRNAs where sequenced using a modified YAMAT-seq (16) protocol previously described in (65). Detailed library methods can be found in Supplementary Methods. Briefly, total RNA was extracted using TRIzol. Total RNA was then deacylated (100 mM Tris-HCl, pH 9.0, 37 °C, 60 min), ethanol precipitated and ligated to YAMAT adapters using T4 RNA ligase 2 (New England Biolabs). Reverse transcription was performed with SuperScript IV (Invitrogen). To remove free adapters, cDNA was size selected for products between 90-200 bp on the BluePippin (Sage Science). Libraries were amplified by 7 cycles of PCR and sequenced on an Illumina NovaSeq X Plus.

### YAMAT-seq read mapping, differential abundance analysis, and tRNA modification index

All YAMAT-seq data processing scripts can be found at https://github.com/warrenjessica/Mealybug_mirror_mt-tRNAs/

Reference database for mapping: The nuclear *P. citri* tRNA reference was generated by scanning the Wellcome nuclear assembly (GCA_950023065) with tRNAscan-SE using the general search option (-G). Mt-tRNA references were determined previously (see annotation section above). The endosymbiont tRNA databases were based on previously published assemblies (CP002244 and NC_015735). Combined tRNA database can be found at (Supp. Table 4).

Read mapping: Reads were either mapped globally to the mitogenome using BLAST (blastn -dust no) to measure detection of tRNA reads from all positions on the mitogenome, or reads were mapped to the combined nuclear, mitochondrial, and endosymbiont tRNA database using a previously described pipeline (65). Differential abundance analysis: Three YAMAT-seq libraries were made from gradient-purified mitochondria, and three YAMAT-seq libraries were made from total insect RNA (three whole insects were homogenized in TRIzol per replicate). Differential abundance analysis was then performed on mapped read counts using EdgeR (66) (Supp. table 6).

tRNA modification index: Generation of a tRNA modification index was done with a pre-existing pipeline (65). A tRNA position was considered modified if >20% of the mapped reads to the reference had a misincorporation at this position.

### Aminoacylation assay

To measure tRNA expression and infer aminoacylation To measure tRNA expression and infer aminoacylation states based on retention of CCA tails, an MSR-seq protocol (30) was performed on three biological replicates from three gradient-purified mitochondrial samples, both with and without periodate treatment. Library construction, sequencing, and data analysis were performed as described previously (43), except that a series of lowered PCR cycles was performed (6 and 10 cycles) in addition to the 15 cycles used in the published protocol. The reference tRNA set for mapping included combined *P*.*citri* database (Supp. Table 4) and a Bacillus subtilis tRNA-Ile that was synthesized and used as an internal “spike-in” control as described previously (43).

### Mitoribosome tRNA-seq

Mitochondrial ribosomes were isolated from crudely isolated mitochondria. Detailed mitoribosome tRNA-seq methods can be found in Supplementary Methods. Briefly, mitochondria were homogenized in lysis buffer containing 25 mM HEPES-KOH, pH 7.4, 150 mM KCl, 50 mM MgOAc, 1.5% β-DDM, 0.15 mg/ml Cardiolipin, 2 mM DTT, 10 mM chloramphenicol, 10 mM cycloheximide, and Pierce Protease inhibitor. The homogenate was spun at 25,000g for 20 min at 4°C in a fixed angle rotor to remove insoluble material. Supernatant was then layered on top of a 2.5 mL 1M sucrose cushion (1M sucrose, 20 mM HEPES, 100 mM KCl, 20 mM MgOAc, 0.6% B-DDM, 0.06 mg/ml Cardiolipin, 2 mM DTT). The sample was centrifuged for 230,000g for 90 min at 4 °C. The supernatant was discarded, and the pellet was carefully washed with resuspension (20 mM HEPES-KOH, pH 7.4,100 mM KCl, 10 mM MgOAc, 0.15% β-DDM, 0.015 mg/ml Cardiolipin, 2 mM DTT) buffer to remove residual sucrose. Ribosome pellet was gently resuspended in resuspension buffer then layered on top of a 10-40% continuous sucrose gradient (in 20 mM HEPES-KOH, pH 7.4, 100 mM KCl, 10 mM MgOAc, 2 mM DTT, created with the BioComp Gradient Master) and spun at 200,000g for 200 min at 4 °C in a swinging bucket rotor. Gradients were fractionated using a BioComp Piston Gradient Fractionator while measuring the 260 nm absorption to identify ribosome fractions. Fractions containing mitoribosomes were pooled, and RNA was extracted using TRIzol using manufacturer’s RNA extraction protocol. YAMAT libraries were then constructed from the resulting RNA.

### PacBio IsoSeq

Isoseq read processing: IsoSeq libraries were created from two biological replicates of total insect RNA and two biological replicates of dissected bacteriome tissue (specialized organ tissue). Libraries were constructed with the SMRTbell prep kit 3.0 (v02.2022) and sequenced on the PacBio Revio instrument. Adapters and poly-A tails were trimmed using cutadapt 5.1 to use the --poly-a option.

IsoSeq read mapping and ORF searches: All trimmed IsoSeq reads were BLASTed (blastn -dust no -evalue 1e-6, query_coverage > 90, percent_identity > 90) to the *P*.*citri* mitogenome assembly to find high confidence mitochondrial reads. These reads were then searched for all ORFs using the EMBOSS: getorf program (-table 5 -find 1 -minsize 300). All ORFS were then BLASTed to the P. manihoti protein coding sequences (tblastn -evalue 1e-6 -seg no) to find homologous ORFs.

### Proteomics

For detailed proteomics methods see Supplementary Methods. Peptides from gradient-purified P. citri mitochondria were analyzed by LC-MS/MS on an Orbitrap Fusion Lumos mass spectrometer coupled to nanoLC. Data were acquired in data-dependent acquisition mode with Orbitrap detection for MS1 and MS2 scans. Raw files were processed using Proteome Discoverer (v3.1).

## Acknowledgments

We thank Courtney York for the mealybug husbandry. This work was supported by the Howard Hughes Medical Institute and a grant from the National Science Foundation (MCB-2322154).

## Data availability

All YAMAT- and MSR-seq reads were deposited on Genbank BioProject (Accession PRJNA1356551). *P. citri* mitogenome assembly GenBank: PZ166947. Scripts used for analysis can be found at https://github.com/warrenjessica/Mealybug_mirror_mt-tRNAs

## Works cited

1. C. Saccone, et al., Mitochondrial DNA in metazoa: degree of freedom in a frozen event. Gene 286, 3–12 (2002).

2. M. A. T. Rubio, et al., Mammalian mitochondria have the innate ability to import tRNAs by a mechanism distinct from protein import. Proc. Natl. Acad. Sci. U.S.A. 105, 9186–9191 (2008).

3. K. Watanabe, Unique features of animal mitochondrial translation systems. The non-universal genetic code, unusual features of the translational apparatus and their relevance to human mitochondrial diseases. Proc Jpn Acad Ser B Phys Biol Sci 86, 11–39 (2010).

4. A. Schneider, Mitochondrial tRNA import and its consequences for mitochondrial translation. Annu Rev Biochem 80, 1033–1053 (2011).

5. T. Salinas-Giegé, R. Giegé, P. Giegé, tRNA biology in mitochondria. Int J Mol Sci 16, 4518–4559 (2015).

6. M. Neiman, D. R. Taylor, The causes of mutation accumulation in mitochondrial genomes. Proc Biol Sci 276, 1201–1209 (2009).

7. M. Lynch, Mutation accumulation in transfer RNAs: molecular evidence for Muller’s ratchet in mitochondrial genomes. Mol Biol Evol 13, 209–220 (1996).

8. M. Lynch, Mutation accumulation in nuclear, organelle, and prokaryotic transfer RNA genes. Molecular Biology and Evolution 14, 914–925 (1997).

9. B. Kuhle, J. Chihade, P. Schimmel, Relaxed sequence constraints favor mutational freedom in idiosyncratic metazoan mitochondrial tRNAs. Nat Commun 11, 969 (2020).

10. F. Jühling, J. Pütz, C. Florentz, P. F. Stadler, Armless mitochondrial tRNAs in Enoplea (Nematoda). RNA Biol 9, 1161–1166 (2012).

11. J. Pons, P. Bover, L. Bidegaray-Batista, M. A. Arnedo, Armless mitochondrial tRNAs conserved for over 30 millions of years in spiders. BMC Genomics 20, 665 (2019).

12. J. M. Warren, D. B. Sloan, Hopeful monsters: unintended sequencing of famously malformed mite mitochondrial tRNAs reveals widespread expression and processing of sense–antisense pairs. NAR Genomics and Bioinformatics 3, lqaa111 (2021).

13. A. Butenko, J. Lukeš, D. Speijer, J. G. Wideman, Mitochondrial genomes revisited: why do different lineages retain different genes? BMC Biol 22, 15 (2024).

14. P. P. Chan, B. Y. Lin, A. J. Mak, T. M. Lowe, tRNAscan-SE 2.0: improved detection and functional classification of transfer RNA genes. Nucleic Acids Research 49, 9077–9096 (2021).

15. M. Bernt, et al., MITOS: Improved de novo metazoan mitochondrial genome annotation. Molecular Phylogenetics and Evolution 69, 313–319 (2013).

16. M. Shigematsu, et al., YAMAT-seq: an efficient method for high-throughput sequencing of mature transfer RNAs. Nucleic Acids Res 45, e70 (2017).

17. J. M. Warren, et al., Combining tRNA sequencing methods to characterize plant tRNA expression and post-transcriptional modification. RNA Biol 18, 64–78 (2021).

18. K. Watanabe, S.-I. Yokobori, tRNA Modification and Genetic Code Variations in Animal Mitochondria. J Nucleic Acids 2011, 623095 (2011).

19. J. M. Warren, et al., Rapid Shifts in Mitochondrial tRNA Import in a Plant Lineage with Extensive Mitochondrial tRNA Gene Loss. Mol Biol Evol 38, 5735–5751 (2021).

20. R. Lorenz, et al., ViennaRNA Package 2.0. Algorithms Mol Biol 6, 26 (2011).

21. T. Suzuki, The expanding world of tRNA modifications and their disease relevance. Nat Rev Mol Cell Biol 22, 375–392 (2021).

22. S. K. Schultz, et al., Modifications in the T arm of tRNA globally determine tRNA maturation, function, and cellular fitness. Proc. Natl. Acad. Sci. U.S.A. 121, e2401154121 (2024).

23. W. C. Clark, M. E. Evans, D. Dominissini, G. Zheng, T. Pan, tRNA base methylation identification and quantification via high-throughput sequencing. RNA 22, 1771–1784 (2016).

24. V. Potapov, et al., Base modifications affecting RNA polymerase and reverse transcriptase fidelity. Nucleic Acids Res 46, 5753–5763 (2018).

25. R. Hauenschild, et al., The reverse transcription signature of N-1-methyladenosine in RNA-Seq is sequence dependent. Nucleic Acids Res 43, 9950–9964 (2015).

26. K. Katoh, J. Rozewicki, K. D. Yamada, MAFFT online service: multiple sequence alignment, interactive sequence choice and visualization. Brief Bioinform 20, 1160–1166 (2019).

27. T. Suzuki, et al., Complete chemical structures of human mitochondrial tRNAs. Nat Commun 11, 4269 (2020).

28. Y. Nakano, et al., Genome-wide profiling of tRNA modifications by Induro-tRNAseq reveals coordinated changes. Nat Commun 16, 1047 (2025).

29. D. V. Lavrov, W. M. Brown, J. L. Boore, A novel type of RNA editing occurs in the mitochondrial tRNAs of the centipede Lithobius forficatus. Proc Natl Acad Sci U S A 97, 13738–13742 (2000).

30. C. P. Watkins, W. Zhang, A. C. Wylder, C. D. Katanski, T. Pan, A multiplex platform for small RNA sequencing elucidates multifaceted tRNA stress response and translational regulation. Nat Commun 13, 2491 (2022).

31. Kristian Davidsen, Lucas B Sullivan (2024) A robust method for measuring aminoacylation through tRNA-Seq eLife 12:RP91554

32. I. Kozarewa, et al., Amplification-free Illumina sequencing-library preparation facilitates improved mapping and assembly of (G+C)-biased genomes. Nat Methods 6, 291–295 (2009).

33. S. Aibara, J. Andréll, V. Singh, A. Amunts, Rapid Isolation of the Mitoribosome from HEK Cells. J Vis Exp 57877 (2018). 10.3791/57877.

34. A. Brown, et al., Structure of the large ribosomal subunit from human mitochondria. Science 346, 718–722 (2014).

35. J. P. Adelman, C. T. Bond, J. Douglass, E. Herbert, Two Mammalian Genes Transcribed from Opposite Strands of the Same DNA Locus. Science 235, 1514–1517 (1987).

36. Y. A. Khan, et al., Evidence for a novel overlapping coding sequence in POLG initiated at a CUG start codon. (2020). 10.17863/CAM.56759.

37. V. Pelechano, L. M. Steinmetz, Gene regulation by antisense transcription. Nat Rev Genet 14, 880–893 (2013).

38. J. Ferrer, N. Dimitrova, Transcription regulation by long non-coding RNAs: mechanisms and disease relevance. Nat Rev Mol Cell Biol 25, 396–415 (2024).

39. A. R. D’Souza, M. Minczuk, Mitochondrial transcription and traslation: overview. Essays Biochem 62, 309–320 (2018).

40. Z. Pietras, et al., Controlling the mitochondrial antisense - role of the SUV3-PNPase complex and its co-factor GRSF1 in mitochondrial RNA surveillance. Mol Cell Oncol 5, e1516452 (2018).

41. G. Santonoceto, A. Jurkiewicz, R. J. Szczesny, RNA degradation in human mitochondria: the journey is not finished. Hum Mol Genet 33, R26–R33 (2024).

42. K. Pawar, et al., Exploration of CCA-added RNAs revealed the expression of mitochondrial non-coding RNAs regulated by CCA-adding enzyme. RNA Biol 16, 1817–1825 (2019).

43. L. F. Ceriotti, J. M. Warren, M. V. Sanchez-Puerta, D. B. Sloan, The landscape of Arabidopsis tRNA aminoacylation. Plant J 120, 2784–2802 (2024).

44. I. Ozerova, et al., Aberrant Mitochondrial tRNA Genes Appear Frequently in Animal Evolution. Genome Biol Evol 16, evae232 (2024).

45. H. Schü rer, S. Schiffer, A. Marchfelder, M. Mörl, This Is the End: Processing, Editing and Repair at the tRNA 3-Terminus. Biological Chemistry 382 (2001).

46. K. Tomita, T. Ueda, K. Watanabe, RNA Editing in the Acceptor Stem of Squid Mitochondrial tRNATyr. Nucleic Acids Research 24, 4987–4991 (1996).

47. A. Reichert, U. Rothbauer, M. Mörl, Processing and Editing of Overlapping tRNAs in Human Mitochondria. Journal of Biological Chemistry 273, 31977–31984 (1998).

48. V. Doublet, et al., Large gene overlaps and tRNA processing in the compact mitochondrial genome of the crustacean Armadillidium vulgare. RNA Biol 12, 1159–1168 (2015).

49. V. Doublet, C. Souty-Grosset, D. Bouchon, R. Cordaux, I. Marcadé, A thirty million year-old inherited heteroplasmy. PLoS One 3, e2938 (2008).

50. Z. Chrzanowska-Lightowlers, J. Rorbach, M. Minczuk, Human mitochondrial ribosomes can switch structural tRNAs - but when and why? RNA Biol 14, 1668–1671 (2017).

51. S. E. Masta, The Complete Mitochondrial Genome Sequence of the Spider Habronattus oregonensis Reveals Rearranged and Extremely Truncated tRNAs. Molecular Biology and Evolution 21, 893–902 (2004).

52. P. B. Klimov, B. M. Oconnor, Improved tRNA prediction in the American house dust mite reveals widespread occurrence of extremely short minimal tRNAs in acariform mites. BMC Genomics 10, 598 (2009).

53. X.-F. Xue, W. Deng, S.-X. Qu, X.-Y. Hong, R. Shao, The mitochondrial genomes of sarcoptiform mites: are any transfer RNA genes really lost? BMC Genomics 19, 466 (2018).

54. D. R. Wolstenholme, J. L. Macfarlane, R. Okimoto, D. O. Clary, J. A. Wahleithner, Bizarre tRNAs inferred from DNA sequences of mitochondrial genomes of nematode worms. Proc. Natl. Acad. Sci. U.S.A. 84, 1324–1328 (1987).

55. R. Okimoto, D. R. Wolstenholme, A set of tRNAs that lack either the T psi C arm or the dihydrouridine arm: towards a minimal tRNA adaptor. The EMBO Journal 9, 3405–3411 (1990).

56. H. Seligmann, Undetected antisense tRNAs in mitochondrial genomes? Biol Direct 5, 39 (2010).

57. A. S. Rodin, E. Szathmáry, S. N. Rodin, One ancestor for two codes viewed from the perspective of two complementary modes of tRNA aminoacylation. Biol Direct 4, 4 (2009).

58. S. N. Rodin, S. Ohno, Two types of aminoacyl-tRNA synthetases could be originally encoded by complementary strands of the same nucleic acid. Orig Life Evol Biosph 25, 565–589 (1995).

59. M. Lynch, The origins of genome architecture (Sinauer Associates, 2007).

60. D. M. Rand, Mitigating Mutational Meltdown in Mammalian Mitochondria. PLoS Biol 6, e35 (2008).

61. C.-H. Kuo, N. A. Moran, H. Ochman, The consequences of genetic drift for bacterial genome complexity. Genome Res 19, 1450–1454 (2009).

62. K. Tamura, Origins and Early Evolution of the tRNA Molecule. Life (Basel) 5, 1687–1699 (2015).

63. R. Root-Bernstein, Y. Kim, A. Sanjay, Z. F. Burton, tRNA evolution from the proto-tRNA minihelix world. Transcription 7, 153–163 (2016).

64. B. Langmead, S. L. Salzberg, Fast gapped-read alignment with Bowtie 2. Nat Methods 9, 357–359 (2012).

65. J. M. Warren, et al., Combining tRNA sequencing methods to characterize plant tRNA expression and post-transcriptional modification. RNA Biol 18, 64–78 (2021).

66. Y. Chen, L. Chen, A. T. L. Lun, P. L. Baldoni, G. K. Smyth, edgeR v4: powerful differential analysis of sequencing data with expanded functionality and improved support for small counts and larger datasets. Nucleic Acids Res 53, gkaf018 (2025).

67. K. Darty, A. Denise, Y. Ponty, VARNA: Interactive drawing and editing of the RNA secondary structure. Bioinformatics 25, 1974–1975 (2009).

