## Supplementary Methods for "Expression of four mitochondrial tRNAs from only two loci"

### Crude isolation of *P. citri* mitochondria

Mealybugs were reared on small potatoes in a chamber set to 12 hr light/ 12 hr dark at 26°C with 50% relative humidity.

*Sample preparation*

Mealybugs were removed from potatoes, washed in 70% ethanol, and rinsed in ice-cold isolation buffer (50 mM HEPES-KOH pH 7.5, 10 mM KCl, 1.5 mM MgOAc, 70 mM sucrose, 210 mM mannitol, 1 mM EDTA, 1 mM EGTA, 1 mM DTT).

*Homogenization*

Cleaned mealybugs were transferred to a 7 mL glass Dounce homogenizer containing 4 mL fresh isolation buffer and homogenized with 10 strokes using a loose pestle followed by 10 strokes with a tight pestle.

*Differential centrifugation and wax removal*

The homogenate was filtered through four layers of pre-wet cheesecloth and one layer of Miracloth. The filtrate was centrifuged at 1,000 × g for 5 min at 4 °C (fixed-angle rotor, Beckman Ti 90), and the pellet was discarded. The supernatant was subjected to a second 1,000 × g centrifugation for 5 min at 4 °C and the resulting supernatant was filtered through a 100 μm mesh filter. To remove residual wax, the filtrate was passed through a second 100 μm mesh filter.

*Mitochondrial pelleting*

The clarified supernatant was centrifuged at 15,000 × g for 15 min at 4 °C to pellet crude mitochondria. The supernatant was discarded.

*Resuspension and downstream use*

The mitochondrial pellet was resuspended in 1 mL SEM buffer (250 mM sucrose, 20 mM HEPES-KOH pH 7.5, 1 mM EDTA, 1 mM EGTA) using a fine paintbrush. Crude mitochondrial preparations were either used immediately for sucrose gradient purification (see below) or snap-frozen in liquid nitrogen for subsequent ribosome isolation or DNA extraction.

### Gradient purification of *P. citri* mitochondria

Resuspended crude mitochondrial pellet was transferred to a 2 mL glass dounce and dounced 4x times with the loose pestle. This suspension was carefully layered on top of a 15-23-32-60% sucrose gradient (in 50mM HEPES-KOH pH7.5, 1 mM EDTA, 1mM EGTA) in a swinging bucket rotor (Beckman Ti 40). The gradient was spun at 28,000g for 1 hr at 4 °C. Purified mitochondria formed a band at the 32/60% interface and was collected by aspiration. Purified mitochondria were washed by adding 5X volumes of SEM buffer and spun at 15,000g for 15 min. Supernatant was discarded and the pellet was snap frozen in liquid nitrogen for further analyses.

Full-length tRNA sequencing

*RNA extraction*

Total RNA for tRNA sequencing was extracted using 1 mL TRIzol following TRIzol manufacturer's RNA extraction protocol.

*Deacylation*

For each YAMAT-seq library, 5 µg total RNA was deacylated in 100 mM Tris-pH 9.0 at 37°C for 60 min. This reaction was neutralized by addition of sodium acetate pH 4.8 to a final concentration of 180 mM. Deacylated RNA was ethanol precipitated and resuspended in RNase free $H_2O$.

*Adapter ligation*

A 9 µl reaction volume containing 1 µg of deacylated RNA and 1 pmol of each Y-5′ adapter (5 pmols total) and 5 pmols of the Y-3′ adapter was incubated at 90°C for 2 min. A total of 1 µl of an annealing buffer containing 500 mM Tris–HCl (pH 8.0) and 100 mM $MgCl_2$ was added to the reaction mixture and incubated for 15 min at 37°C. Ligation was performed by adding one unit of T4 RNA Ligase 2 enzyme (New England Biolabs) in 10 µl of 1× reaction buffer and incubating the reaction at 37°C for 60 min then incubating at 4°C at 60 min. All adapter and primer sequences can be found in Supp. Table 9.

*Reverse transcription*

RT of ligated RNA was performed using SuperScript IV (Invitrogen) according to the manufacturer's protocol but with slight modifications. Briefly, 2 µl of 2 µM RT primer, and 2 µl of 10 mM dNTP mix was added to 20 µl of the ligated RNA. The mixture was briefly vortexed, centrifuged and incubated at 70°C for 2 min and the temperature was reduced to 37 °C by 0.1 °C/s. Then, 8 µl of 5× SSIV buffer, 2 µl 100 mM DTT, 2 µl RNaseOUT and 2 µl of SuperScriptIV was added to each reaction. The reaction was incubated for 10 min at 55 °C and then the samples incubated at 35 °C overnight for an overnight RT reaction.

*Size selection of the cDNA*

To remove free adapters and adapter dimers, a size selection was done on the cDNA products prior to PCR. cDNA was size selected on the BluePippin (Sage Science), using 3% agarose gel cassettes and marker Q3 following the manufacturer's protocol. The size selection parameters were set to a range of 90-200 bp. Size-selected cDNA was then cleaned using solid phase reversible immobilization beads (1.8X ratio) and resuspended in 10 mM Tris-pH 8.0.

*PCR*

The resulting cDNA was amplified by polymerase chain reaction (PCR) in a 50 µl reaction containing 20 µl of the size selected cDNA, 25 µl of the NEBNext 2× PCR Master Mix, 2.5 µl of the PCR forward primer, 2.5 µl of the PCR reverse primer. Seven cycles of PCR were performed with an initial 1 min incubation at 98°C and 7 cycles of 30 s at 98°C, 30 s at 60°C and 30 s at 72°C, followed by 5 min at 72°C. PCR products were cleaned using solid phase reversible immobilization beads (1.6X ratio) and resuspended in 10 mM Tris (pH 8.0). Libraries were sequenced on an Illumina NovaSeq X Plus with paired-end, 150-bp reads.

### Mitoribosome tRNA-seq

*Mitochondrial ribosome isolation*

Mitochondrial ribosomes were isolated from crudely isolated mitochondria. Each mitoribosome isolation used three pooled, crude mitochondrial isolations, totaling approximately 6 g of mealybugs. Mitochondria were resuspended in 1.5 mL of lysis buffer containing 25 mM HEPES-KOH, pH 7.4, 150 mM KCl, 50 mM MgOAc, 1.5% β-DDM, 0.15 mg/ml Cardiolipin, 2 mM DTT, 10 mM chloramphenicol, 10 mM cycloheximide, and a Pierce Protease Inhibitor Mini Tablets, EDTA-free (1 tablet/10 ml). Solution was transferred to a 2 mL glass dounce and dounced 20x with the tight pestle avoiding the generation of bubbles. Homogenized mitochondria were placed on a roller for 30 min at 4°C to complete mitochondrial lysis. The homogenate was spun at 25,000g for 20 min at 4°C in a fixed angle rotor to remove insoluble material. Pellet was discarded by carefully decanting the supernatant. Supernatant was then layered on top of a 2.5 mL 1M sucrose cushion (1M sucrose, 20 mM HEPES, 100 mM KCl, 20 mM MgOAc, 0.6% B-DDM, 0.06 mg/ml Cardiolipin, 2 mM DTT). The sample was centrifuged for 230,000g for 90 min at 4 °C. The supernatant was discarded, and the pellet was carefully washed with resuspension (20 mM HEPES-KOH, pH 7.4,100 mM KCl, 10 mM MgOAc, 0.15% β-DDM, 0.015 mg/ml Cardiolipin, 2 mM DTT) buffer to remove residual sucrose. Ribosome pellet was gently resuspended in a fresh 300 µl resuspension buffer by rocking back and forth, using vortex on a very low speed, and gentle pipetting. The resuspended ribosomes were then layered on top of a 10-40% continuous sucrose gradient (in 20 mM HEPES-KOH, pH 7.4, 100 mM KCl, 10 mM MgOAc, 2 mM DTT, created with the BioComp Gradient Master) and spun in a Beckman Ti 40 rotor at 200,000g for 200 min at 4 °C. Gradients were fractionated using a BioComp Piston Gradient Fractionator while measuring the 260 nm absorption to identify ribosome fractions. Fractions containing mitoribosomes were pooled, and RNA was extracted using TRIzol using manufacturer's RNA extraction protocol.

YAMAT libraries were then constructed from the resulting RNA.

*Identification of ribosomes in sucrose gradient fractions*

Total RNA was extracted from the sucrose gradient fractions 5-12 separately and RT-qPCR was used to determine the relative abundance of each ribosome type in the gradient. 100 ng of RNA from each fraction was reverse transcribed (iScript Reverse Transcription Supermix, Bio-rad) using manufacturer's protocol. 1 µl of the cDNA reaction mixture for each fraction was then subjected to qPCR targeting primers for the large ribosomal rRNA from the cytosolic, mitochondrial, and endosymbiont ribosomes (primers can be found in Supp. Table 10B).

### PacBio IsoSeq

*Isoseq read processing*

IsoSeq libraries were created from two biological replicates of total insect RNA and two biological replicates of dissected bacteriome tissue (specialized organ tissue). Libraries were constructed with the SMRTbell prep kit 3.0 (v02.2022) and sequenced on the PacBio Revio instrument. Adapters and poly-A tails were trimmed using cutadapt 5.1 to use the --poly-a option.

*IsoSeq read mapping and ORF searches*

All trimmed IsoSeq reads were BLASTed (blastn -dust no -evalue 1e-6, query_coverage > 90, percent_identity > 90) to the *P.citri* mitogenome assembly to find high confidence mitochondrial reads. These reads were then searched for all ORFs using the EMBOSS: getorf program (-table 5 -find 1 -

minsize 300). All ORFS were then BLASTed to the *P. manihoti* protein coding sequences (tblastn -evalue 1e-6 -seg no) to find homologous ORFs.

### Proteomics

Peptides were isolated and sequenced from gradient-purified *P. citri* mitochondria at the Arizona State University Mass Spectrometry Facility. Peptides were injected onto an Ultimate 3000 HPLC (Thermo Fisher Scientific) at 0.300 nL/min and separated on a 50 cm EASY SPRAY C18 column (Thermo Fisher Scientific) before being analyzed in an Orbitrap Fusion Lumos Tribrid Mass Spectrometer (Thermo Fisher Scientific). The mass spectrometer was run in database-dependent acquisition (DDA) mode with both MS1 and MS2 being detected on the Orbitrap. The MS1 full scan resolution was set to 120K, a scan range of 375-1500 m/z, charge states were set to the range of +2 to +7, the AGC target was set to 400K, and maximum injection time was 50ms. Precursor ions were activated by HCD with 30% collision energy, MS2 resolution was set to 30K, the AGC target was set to 50K, and the maximum injection time was 54ms. Raw data files were analyzed in Proteome Discoverer 3.1 (Thermo Fisher Scientific).
