## Supplementary Figures for "Expression of four mitochondrial tRNAs from only two loci"

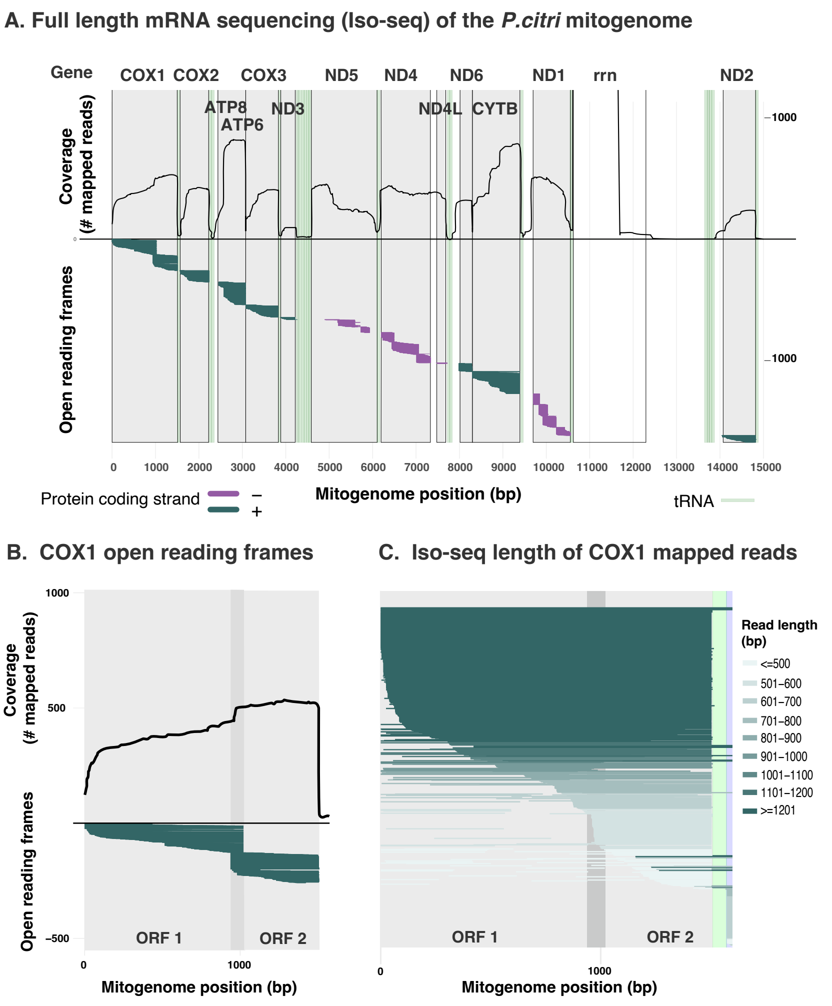


**Supplementary Figure 1| IsoSeq coverage and the resulting ORFs for *P. citri* mitochondrial protein-coding genes.** A. Top row shows the coverage (number of reads) of IsoSeq reads mapped to the mitogenome with the gene name above. The bottom row shows all the predicted ORFs sequences found in the mapped IsoSeq reads that have homology to *P. manihoti* (blastp, disabled low-complexity filtering, e-value < 1e-6). The coding strand of the ORFs are indicated by color, with genes on the positive strand in green, those on the reverse stand in purple. Genomic locations of mt-tRNA gene locations are shown in light green shading. COX1, ND1, ND4, and ND5 do not exist as a single, continuous ORFs, which can be seen from no single ORF spanning the entire region of those genes. B. Detail of the expression level and resulting ORFs of the COX1 gene. The region mapping to the COX1 has two ORFs, with a small amount of overlap (shown in a dark gray stripe). C. IsoSeq reads that map to COX1 region shorted and colored by length. The majority of the IsoSeq reads are greater than 1201 bp and contain both ORFs. Overlap region between the two ORFs is highlighted in a dark gray stripe. Mt-tRNA-Leu(TAA) gene shown in light green stripe immediately to the right of the second COX1 ORF.

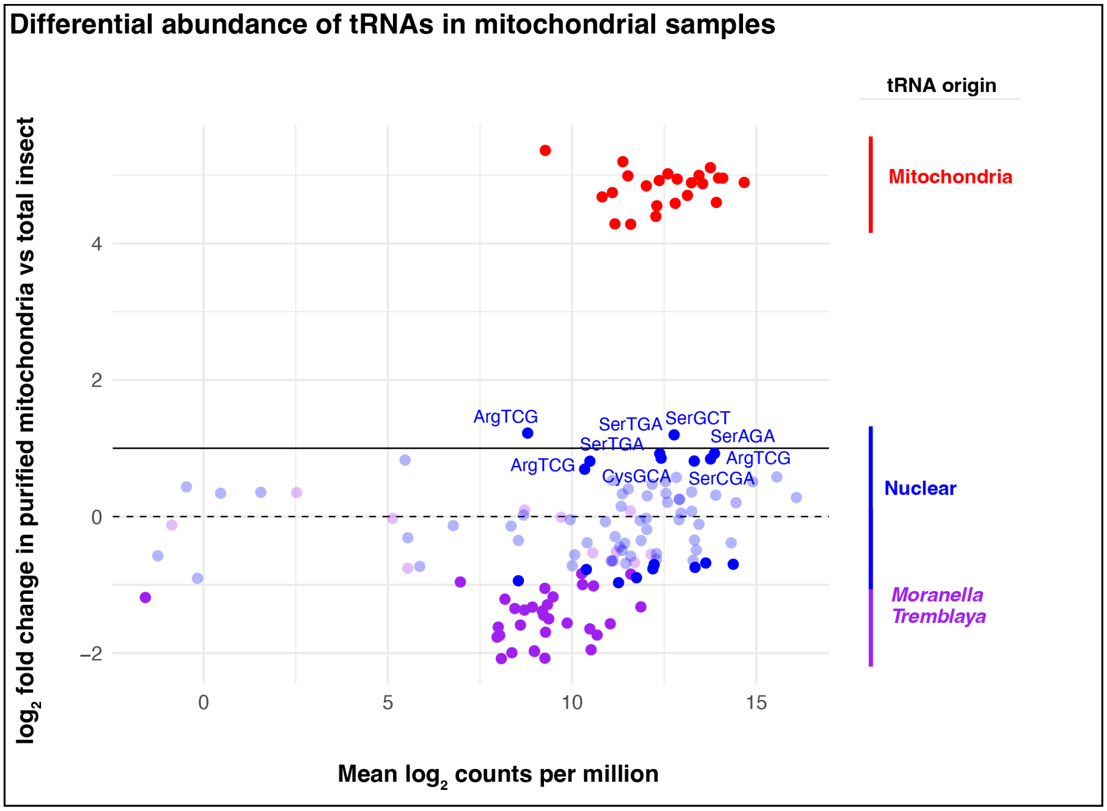


**Supplementary Figure 2| Differential abundance of all *P. citri* tRNAs in gradient-purified mitochondria compared to total insect samples.** Each point represents a unique tRNA reference and is colored by the encoding genome (mitochondrial tRNAs in red, nuclear tRNAs in blue, and both nutritional endosymbionts tRNAs in purple). Darker dots are tRNAs with an FDR value > 0.05. Dashed line is at 0 (lack of enrichment or depletion), and the thick line is at $\log_2$ fold enrichment of 1 (2x). Nuclear tRNAs with a positive enrichment and an FDR significance > 0.05 are also indicated with the amino acid and anticodon.

Folding models of all *P. citri* mt-tRNAs

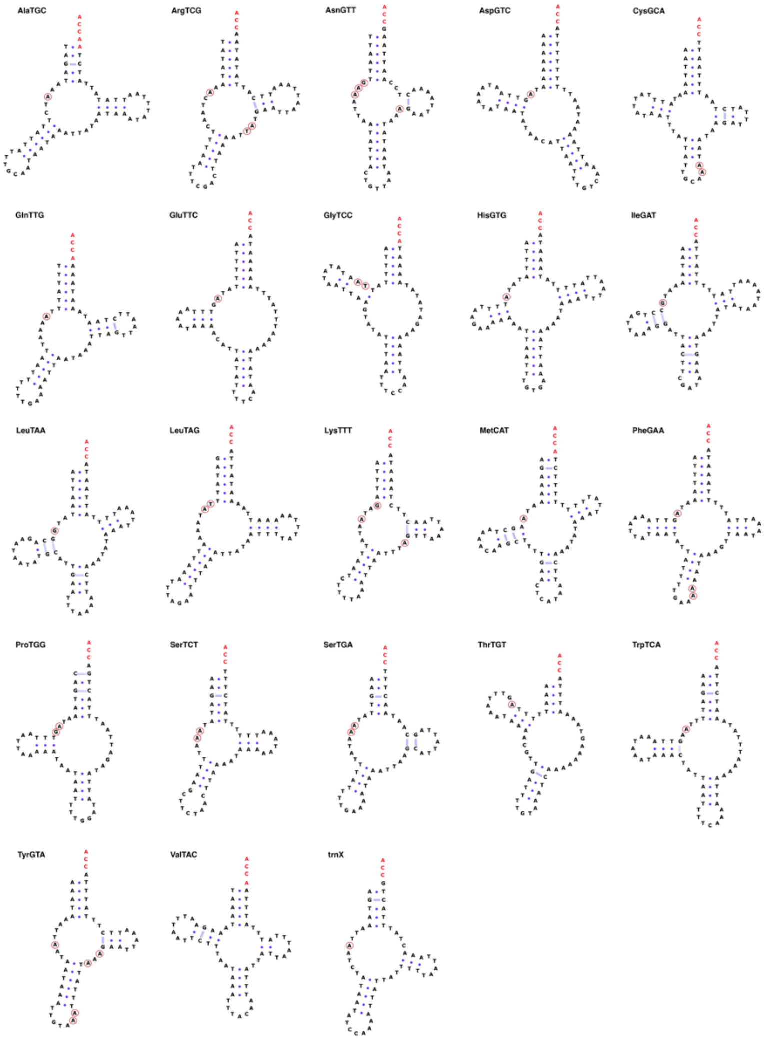


**Supplementary Figure 3| Predicted folded structure diagrams of all *P. citri* mt-tRNA.** Mt-tRNAs were manually folded to center anticodon and prioritize stem and loop structure. Folded structure diagrams were created using VARNA ver. 3.9 (Darty et al. 2009).

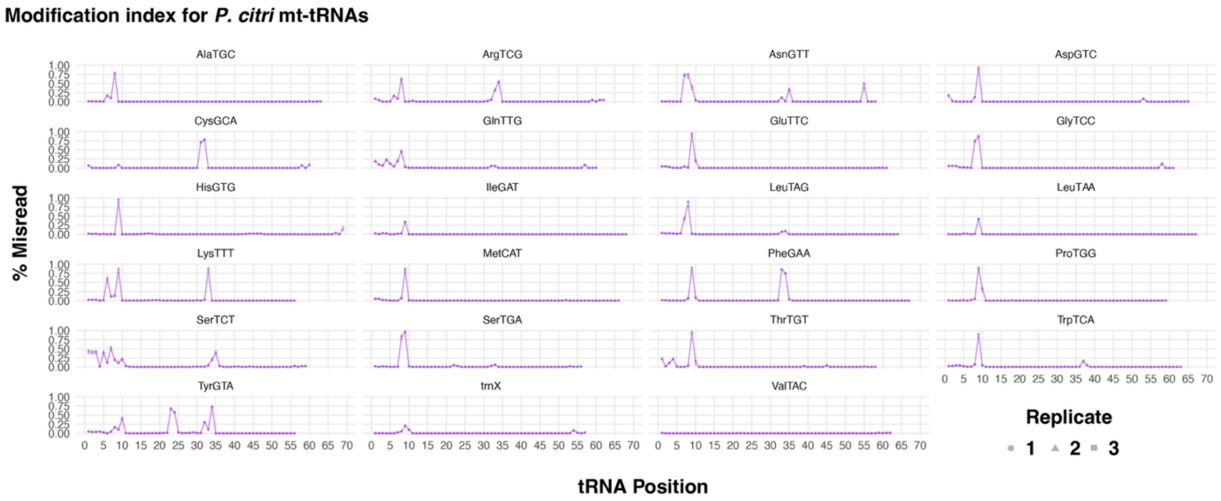


**Supplementary Figure 4| Modification indexes for all mt-tRNAs.** Modification indexes for mt-RNAs in the three mitochondrial isolation YAMAT libraries. A tRNA position was considered modified if >20% of the mapped reads to the reference had a misincorporation at this position. Some tRNAs had slightly overlapping sequences in the mitogenome and the processing of the upstream or downstream tRNA resulted in a truncated sequence that also produced an overhang captured by YAMAT-seq. These sequences produced a variable end (e.g., SerTCT) but were not considered as candidates for base modifications at these positions in Supp. Fig. 3.

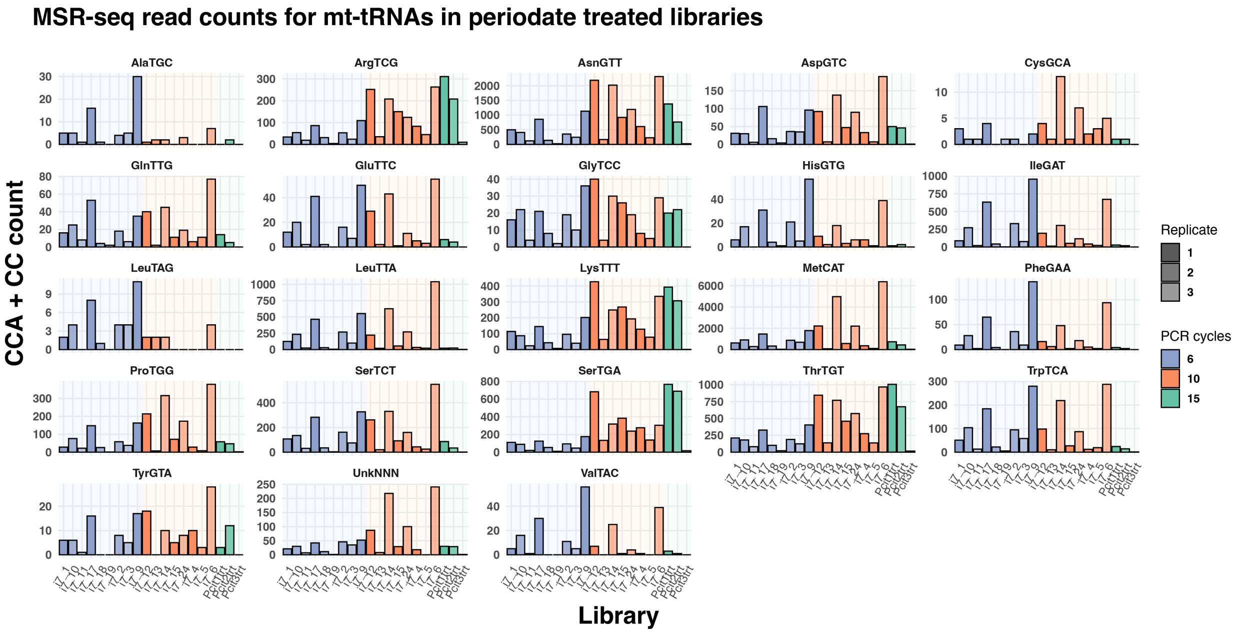


**Supplementary Figure 5| Detection of CCA- and CC-ending reads for each mt-tRNA in MSR-seq libraries.** Each bar is a separate MSR-seq library specified with its unique barcode identifier. Each library was generated from a separate biological replicate, indicated in shading density. The PCR cycle number used in library amplification is indicated by a different color.

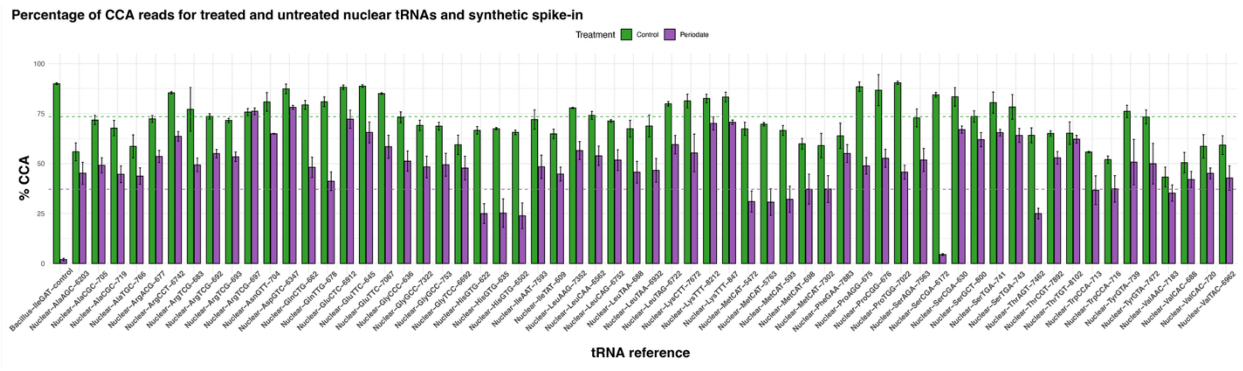


**Supplementary Figure 6| Percentage of reads ending in CCA or CC for nuclear *P. citri* tRNAs.** Only nuclear tRNAs with at least 1,000 reads combined in both the periodate treated and untreated libraries are shown. Total MSR-seq read counts for all references can be found in Supp. Table 7, and all CCA/CC/other read averages for all *P. citri* tRNA reference averages can be found in Supp. Table 8.

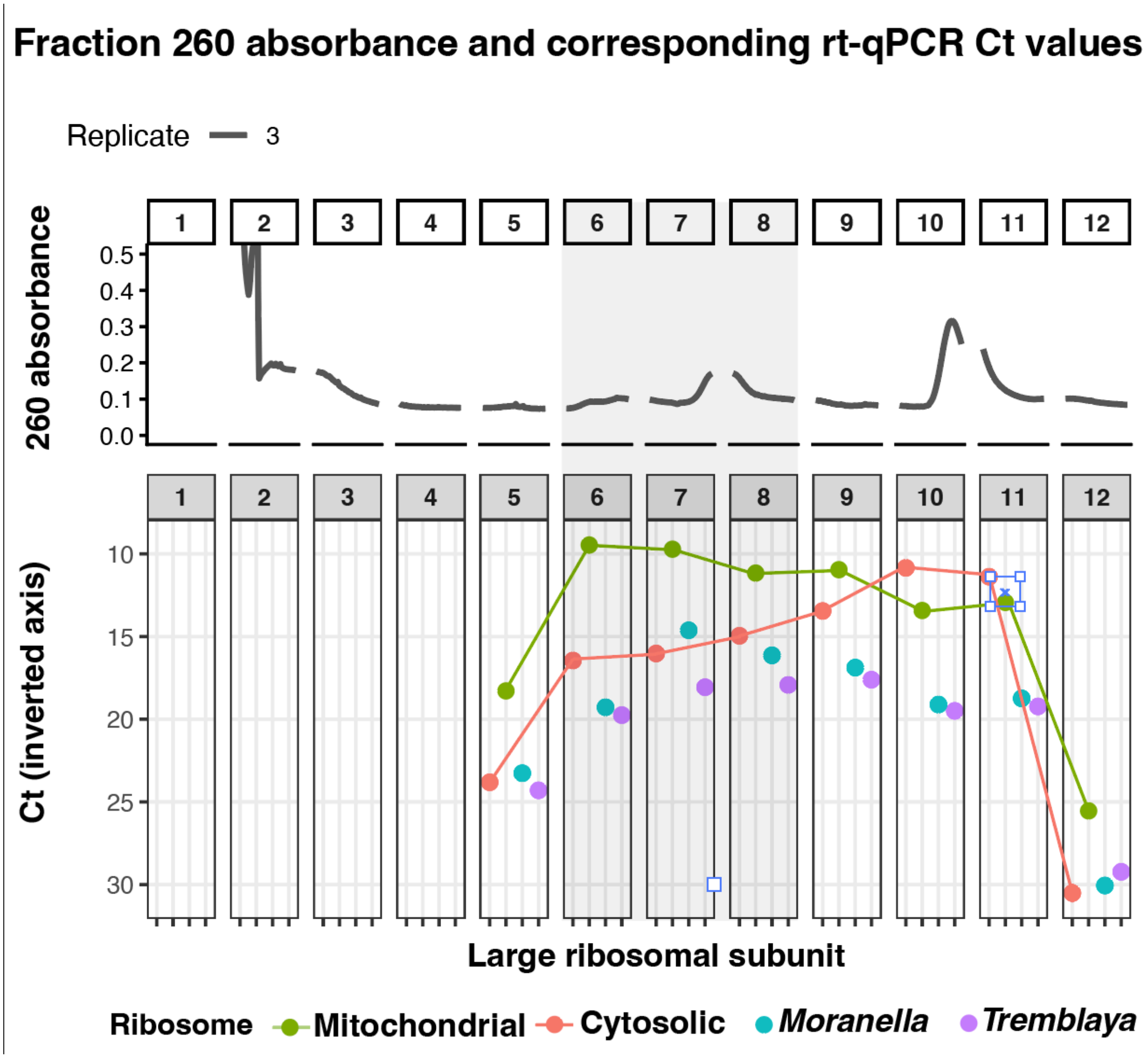


**Supplementary Figure 7| rRNA distribution across sucrose gradient fractions.** 260-nm absorbance profile (top) and corresponding RT-qPCR cycle threshold (Ct) values for large-subunit rRNA markers across sucrose gradient fractions. Ct values (bottom; inverted axis) are shown for mitochondrial, cytosolic, and the two endosymbiont ribosomes (*Moranella* and *Tremblaya*). All samples were normalized to 5 ng of cDNA for qPCR. Lower Ct values indicate higher rRNA abundance in a fraction. Points represent the mean of three qPCR replicates. The 260-nm absorbance trace is shown for experimental replicate 3.
